# Comparative analysis of novel *Pseudobdellovibrionaceae* genera and species yields insights into the genomics and evolution of bacterial predation mode

**DOI:** 10.1101/2025.02.19.638989

**Authors:** Rebecca L. Maher, Janna Wülbern, Johannes Zimmermann, Emily Yeh, Liesl Benda, Urska Repnik, Janina Fuß, Sven Künzel, Hinrich Schulenburg, Brendan J.M. Bohannan, Karen L. Adair, Julia Johnke

**Affiliations:** Friday Harbor Laboratories, University of Washington, Friday Harbor, Washington, USA; Institute of Ecology and Evolution, Department of Biology, University of Oregon, Eugene, Oregon, USA; Evolutionary Ecology and Genetics, Zoological Institute, Kiel University, Kiel, Germany; Cluster of Excellence Balance of the Microverse, Friedrich Schiller University, Jena, Germany; Central Microscopy, Department of Biology, Kiel University, Kiel, Germany; Institute of Clinical Molecular Biology (IKMB), Kiel University, Kiel, Germany; Max Planck Institute for Evolutionary Biology, Plön, Germany; Antibiotic Resistance Group, Max Planck Institute for Evolutionary Biology, Plön, Germany

## Abstract

Bacteria of the family *Pseudobdellovibrionaceae* belong to a group of bacteria that kill and feed on other bacteria. The diversity of predation strategies, habitats, and genome characteristics of these bacteria are largely unexplored, despite their ecological and evolutionary importance in microbial communities. Therefore, we characterized new *Pseudobdellovibrionaceae* strains isolated from the direct environments of three animal hosts: the zebrafish (*Danio rerio*), the threespine stickleback fish (*Gasterosteus aculeatus*), and the nematode *Caenorhabditis elegans*. We used transmission electron microscopy (TEM) and genomic analyses to characterize the morphology and predation modes of our isolates. While most of our isolates exhibited periplasmic (i.e. endoparasitic) predation, one isolate clearly exhibited epibiotic (i.e. exoparasitic) predation and represents only the third confirmed epibiotic strain within its clade. Of our isolates, six are members of five new species in the genus *Bdellovibrio* and two strains likely represent new genera within the family *Pseudobdellovibrionaceae*. From metabarcoding data, we found indications that *Pseudobdellovibrionaceae* are widespread among our three animal hosts. Genomic analyses showed that epibiotic predators lack genes involved in host independence (i.e. prey-independent feeding) and peptidoglycan modification. However, genes unique to epibiotic predators may underlie this predation mode, particularly those involved in cell wall remodeling and recycling. With robust phylogenomic analyses, we show that our novel isolates cluster with previously described *Pseudobdellovibrionaceae* isolates according to predation mode. Further, by placing *Pseudobdellovibrionaceae* predators within a wider evolutionary history including other predatory and non-predatory bacteria, we postulate periplasmic predation as the ancestral mode, with more derived epibiotic predators exhibiting genome streamlining.

## Introduction

For decades, researchers have documented the mechanisms by which top-down control via predation drives structure and diversity in communities of plants and animals [1, 2]. More recent evidence suggests that predation by bacteria may similarly structure microbial communities, including those associated with animal hosts (e.g. in corals [3, 4]). Moreover, the presence, abundance, and richness of bacterial predators are positively correlated with overall microbiome diversity for numerous host-microbiome systems, most likely because predation on highly abundant species allows rare taxa to thrive [5]. Understanding the potential contribution of bacterial predators to host microbiome dynamics is important given the central role that microbiomes can play in host health and well-being.

One of the best-studied bacterial predators, *Bdellovibrio bacteriovorus*, was discovered over 60 years ago [6]. It is now considered a member of the “*Bdellovibrio* and like organisms” (BALOs), a group of obligate predators of Gram-negative bacteria. Since then, detailed information has been obtained on the distribution of BALOs across terrestrial and aquatic habitats and their complex strategies of predation, growth, and reproduction [7, 8]. BALOs play a role in microbial population control and nutrient cycling through substantial contributions to bacterial death, despite their relatively low abundance [9]. However, most BALO research has been restricted to a few type strains, and we are only beginning to uncover and comprehensively describe the taxonomic and genetic diversity of BALOs [10]. For instance, our understanding of bacterial predation modes largely reflects research done with two type strains from the *Pseudobdellovibrionaceae* family (formerly *Bdellovibrionaceae* [11, 12]): the periplasmic predator *B. bacteriovorus* HD100 and the epibiotic predator *Pseudobdellovibrio. exovorus* JSS. Both predators employ a biphasic lifestyle including an attack phase in which the small, motile predator cells actively search for prey cells. Once anchored to the prey cell, a periplasmic predator enters the prey periplasm, forms a *bdelloplast*, and grows by consuming the prey’s cellular components. Then, the predator cell replicates, septates, and lyses the bdelloplast, releasing offspring cells. In contrast, epibiotic predators attach to the prey cell and consume the prey’s cytoplasmic content without intrusion. The epibiotic predator *P. exovorus* then undergoes binary or non-binary fission [13] and produces two to three attack phase daughter cells.

Next-generation sequencing technologies have uncovered genomic signatures of predation among the BALOs [14], and genetic elements that differentiate the periplasmic and epibiotic predation modes [15]. In an initial evaluation, epibiotic predators were deficient in numerous functions due to their smaller genomes compared to the periplasmic predators.

However, the recent genomic characterization of an epibiotic predator placed within the genus *Bdellovibrio* with a larger genome suggests that the genomic diversity in epibiotic predators is under sampled [16].

In order to expand our understanding of BALO taxonomic and genetic diversity, especially in animal hosts, we isolated nine novel *Pseudobdellovibrionaceae* strains from the immediate environments of three important model animal species: the zebrafish (*Danio rerio*), the threespine stickleback fish (*Gasterosteus aculeatus*), and the nematode *Caenorhabditis elegans*. We characterized these strains using microscopy across the life cycle to infer predation mode. We surveyed metabarcoding data from all three animal hosts to infer the prevalence of these novel strains. Finally, we sequenced and assembled the genomes of these *Pseudobdellovibrionaceae* strains *de novo* and compared them to existing genomes of predators of the genera *Bdellovibrio* and *Pseudobdellovibrio*. These analyses expand substantially on previous comparative genomic analyses of predatory bacteria [14, 15, 17, 18], led to the formal description of new genera and species within the family *Pseudobdellovibrionaceae*, and provide the most comprehensive phylogenetic history to date of predation mode, revealing conserved genomic features differentiating these two predation lifestyles.

## Methods

All materials and methods can be found in the supplemental material.

## Results

### Newly obtained *Pseudobdellovibrionaceae* isolates varied in size and predation mode

From the direct environments of three animal hosts, we isolated 9 novel BALO strains (Table 1). Transmission electron microscopy (TEM) after negative staining was employed to confirm predation strategies (Figure 1, Figure S1). The presence of a bdelloplast containing at least one BALO cell was considered an indication of periplasmic predation. Bacteria of all isolates had a curved-rod (vibrioid) shape and all periplasmic strains had thicker flagellar filaments, which likely represent ensheathed flagella [19] and may help periplasmic predators to retract their flagellum during prey invasion [20]. In contrast, the only epibiotic strain had a notably thinner and longer flagellar filament, which suggests that this strain does not have a sheathed flagellum. Moreover, flagellar waves differed between periplasmic predators and the epibiotic *Bdellovampiro gaculeatus* SBM16 strain (see below for justification and a formal description of all newly identified species). In detail, periplasmic predators showed the tapered waves as described for *Bdellovibrio bacterivorus* [21, 22]. In contrast, the flagellum of *Bdellovampiro gaculeatus* SBM16 consisted of homogenous waves with smaller amplitude.

**Figure 1:**
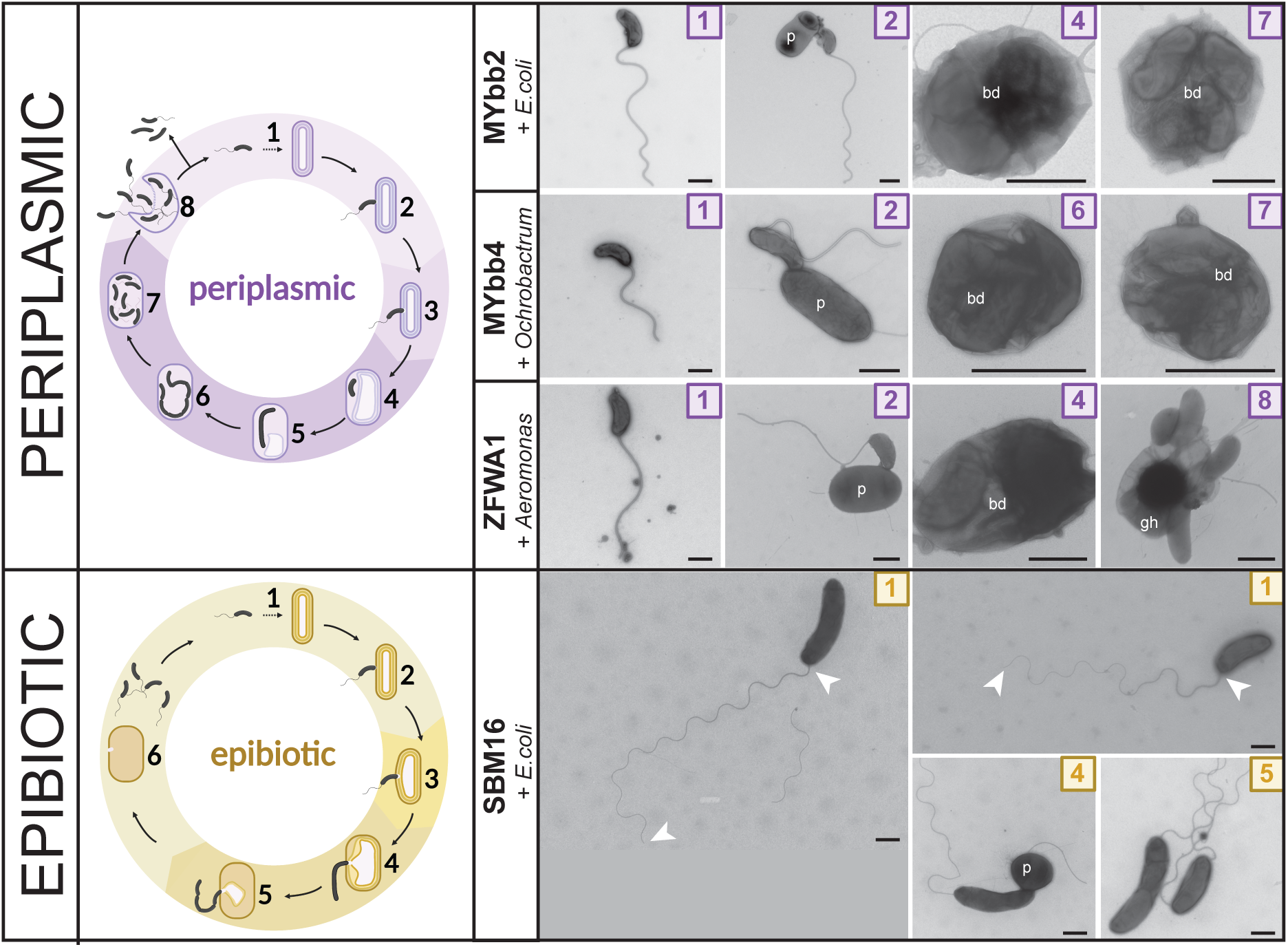
Cell shape and predation mode of *Pseudobdellovibrionaceae* strains MYbb2, MYbb4, ZFWA1 and SBM16. Life cycles of periplasmic (purple) and epibiotic (yellow) *Pseudobdellovibrionaceae* isolates are schematically represented on the left. Numbers in image corners correspond to the proposed life cycle stage. The upper panels illustrate periplasmic *Pseudobdellovibrionaceae* isolates. Stage 1 images represent *Pseudobdellovibrionaceae* cells in attack phase. In stage 2, predators are attached to prey cells (p). Stages 4-7 demonstrate intra- periplasmic growth in bdelloplasts (bd), with one or more *Pseudobdellovibrionaceae* cells confined within the outer membrane of a prey cell. Stage 8 depicts *Pseudobdellovibrionaceae* cells leaving the prey ghost cell (gh). The lower panels display the epibiotic *Bdellovampiro gaculeatus* strain SBM16. This strain has a long flagellum, as illustrated in stage 1 images with white arrows at the start and end point of the flagellum. After attachment to and penetration of the outer membrane, epibiotic cells grow and divide outside prey cells (stages 4-5) as indicated by different sizes of cells and by septation in elongated bacteria. Samples were imaged by TEM after negative staining. Scale bar is 500 nm. Life cycle images created in BioRender. Wülbern, J. (2024) BioRender.com/t31h353.

**Table 1.** Phenotypic characterization of novel *Pseudobdellovibrionaceae* isolates. Values are expressed as mean ± standard error (MYbb1, MYbb2, MYbb4, ZFWA1) or as mean ± standard deviation (MYbb5, MYbb7, MYbb10, MYbb11, SBM16). Respective highest/lowest values are highlighted in bold. Measurements for all independent experiments and grids are given in Table S1.

| Strain | Species<sup>a</sup> | Flagellum length [$\mu\text{m}$] | Cell body area [$\mu\text{m}^2$] | Perimeter [$\mu\text{m}$] | Feret diameter ratio<sup>b</sup> | Cultivation prey | Predation strategy<sup>c</sup> | Morphology<sup>d</sup> |
| --- | --- | --- | --- | --- | --- | --- | --- | --- |
| MYbb11 | *Bdellovibrio bacteriovorus* | 2.87 ± 0.57 | 0.22 ± 0.03 | 1.85 ± 0.17 | 1.81 | *E.coli* | periplasmic | |
| MYbb2 | *Bdellovibrio tiberii* | 3.34 ± 0.36 | 0.25 ± 0.04 | 2.01 ± 0.2 | 1.92 | *E.coli* | periplasmic | |
| MYbb5 | *Bdellovibrio kumpostii* | 3.02 ± 0.55 | 0.21 ± 0.03 | 1.9 ± 0.16 | 2.08 | *E.coli* | periplasmic | narrow cell |
| MYbb1 | *Bdellovibrio tomkyle* | 3.02 ± 0.22 | **0.19** ± 0.02 | **1.67** ± 0.1 | **1.78** | *E.coli* | periplasmic | |
| MYbb7 | *Bdellovibrio bagaluti* | 2.99 ± 0.44 | 0.21 ± 0.03 | 1.82 ± 0.19 | 2.01 | *E.coli* | periplasmic | |
| MYbb10 | *Bdellovibrio bagaluti* | 2.9 ± 0.34 | 0.29 ± 0.04 | 2.36 ± 0.21 | 2.29 | *E.coli* | periplasmic<br>(no confirmation) | narrow cell |
| MYbb4 | *Bdellovibrio krueschi* | **2.58** ± 0.33 | 0.25 ± 0.03 | 2.15 ± 0.2 | 2.16 | *Ochrobactrum sp.* | periplasmic | short flagellum, narrow cell |
| ZFWA1 | *Bdellovibrio daniorerio* | 2.77 ± 0.25 | 0.25 ± 0.05 | 1.99 ± 0.2 | 1.9 | *Aeromonas sp.* | periplasmic | |
| SBM16 | *Bdellovibrio gaculeatus* | **7.13** ± 1.57 | **0.77** ± **0.24** | **4.33** ± **0.97** | **3.24** | *E.coli* | epibiotic | long flagellum, narrow cell |

<sup>a</sup>Species named according to results of comparative genome analysis, see section below for details.
<sup>b</sup>The Feret diameter ratio was used as a proxy for cell size and shape, with higher values indicating elongated cells.
<sup>c</sup>Phenotype confirmed by TEM imaging after negative staining.
<sup>d</sup>Description for strains with outstanding phenotype.

The length of flagella and the size of predator cells varied between different strains (Table 1) and across independent experiments of single strains (Table S1). Cells sampled after the addition of fresh prey appeared larger and had a shorter flagellum than those from prey- cleared standard overnight cultures. Variability in cell size within strains and even within the same culture may be explained by an earlier observation that BALO flagella [23] and cells continue to elongate after exiting the bdelloplast [24]. The negative correlation between flagellum length and cell size is, however, puzzling. Notably, in the epibiotic strain, variation in cell size was larger than in periplasmic strains, as illustrated by two differently sized attack phase cells of SBM16 (Figure 1).

Between strains cell size and flagellum length showed no correlation (Table 1). For example, the epibiotic *Bdellovampiro gaculeatus* strain SBM16 cells exhibited both a large cell size (0.77 ± 0.24 µm^2^) and the longest flagellum (7.13 ± 1.57 µm) among the analyzed strains (Figure 1). In contrast, *Bdellovibrio tomkyle* strain MYbb1, despite having a small cell size (0.19 ± 0.02 µm^2^), had an average flagellum length (3.02 ± 0.22 µm), while *Bdellovibrio bagaluti* strain MYbb10 displayed a comparable flagellum length (2.9 ± 0.34 µm) but a larger cell size (0.29 ± 0.04 µm^2^) (Table 1, Figure S1). Notably, potential damage to the flagella during sample preparation may have affected the measured flagellum lengths, possibly leading to underestimation.

Cell shape was further assessed using the Feret diameter ratio, which indicates the degree to which a particle is stretched or how similar its projected contour is to a circle, with higher values suggesting elongated shape (Table 1). Notably, the highest values were recorded for *Bdellovibrio krueschi* strain MYbb4, *Bdellovibrio kumpostii* strain MYbb5, *Bdellovibrio bagaluti* strain MYbb10, and *Bdellovampiro gaculeatus* strain SBM16, indicating these cells exhibited elongated or narrow morphologies compared to other BALO cells (Table 1).

### Comparative genomic analysis reveals novel *Pseudobdellovibrionaceae* genera and species

To assign species and genus level taxonomic classifications to the novel strains, we used average nucleotide identity (ANI) (>95% [25]) and average amino acid identity (AAI) (>65% [26]). These comparative genomic analyses suggest novel genus and species relationships within the *Pseudobdellovibrionaceae* family (Figure S2; Table S2). Building on previous work [12, 18], AAI values support the existence of four genera within *Pseudobdellovibrionaceae* including existing (*Bdellovibrio*, *Pseudobdellovibrio*) and novel genera (*Bdellovenatio*, *Bdellovampiro*).

Genus *Bdellovenatio* includes the novel strain *B. daniorerio* ZFWA1 which shares <62% AAI with all other genomes. Genus *Bdellovampiro* includes novel and existing strains *B. gaculeatus* SBM16 and *B. qaytius*, respectively, which share an AAI of 79.59%, and each have an AAI <62% with *Pseudobdellovibrio exovorus* JSS. *Bdellovibrio* species were confirmed for strains MYbb10 and MYbb7 (now *B. bagaluti* strains MYbb10 and MYbb7), strains MYbb2 and *B. bacteriovorus Tiberius* (now *B. tiberii* strains MYbb2 and Tiberius), and strains MYbb11 (now *B. bacteriovorus* strain MYbb11), *B. bacteriovorus* 109J, and *B. bacteriovorus* HD100. Further, the strains MYbb1 (now *B. tomkyle* strain MYbb1), MYbb4 (now *B. krueschi* strain MYbb4), and MYbb5 (now *B. kumpostii* strain MYbb5) were identified as separate species. The results also suggest that *B. bacteriovorus* strains SSB218315, kdesi, and W represent separate new species. Notably, ANI values between 85%-95% are comparatively rare and represent a discontinuity in the distribution of ANI values among closely related genomes [27]. A number of genomes had ANI values within this range, including strains of *B. bacteriovorus*, *B. tiberii*, *B. kumpostii*, and *B.* sp SSB218315, strains *B*. sp ZAP7 and *B*. sp KM01, and strains *B*. sp kdesi and *B. tomkyle* MYbb1. While considered “species-like”, these sequence-discrete populations are generally ecologically differentiated and still considered distinct species [27].

### Phylogenomic analysis identifies two distinct clades for periplasmic and epibiotic *Pseudobdellovibrionaceae*

To uncover the ancestral predation mode in predatory bacteria, a tree with representative outgroups including predatory and non-predatory bacteria, as well as one archaeon (Table S3), was generated. The phylogenetic analysis led to the identification of two distinct clades, with one clade containing all predatory bacteria with a periplasmic lifestyle (Figure 2a). *P. exovorus* and the non-predator *Oligoflexus tunisiensis* clustered within this clade as well. The other predatory bacteria with a cooperative or epibiotic predation strategy (*Myxococcus xanthus*, *Micavibrio aeruginosavorus*, and *Vampirovibrio chlorellavorus*) clustered within the second clade. It should be noted that the genus *Micavibrio* is also part of the family *Pseudobdellovibrionaceae*. Our phylogenomic analysis and others [12, 18] indicates that *Micavibrio* is not in close relation to the known isolates of the genera *Bdellovibrio* and *Pseudobdellovibrio* nor to our newly isolated strains and should be therefore re-classified. We are subsequently omitting *Micavibrio* from the following focused analysis and whenever we discuss members of the family *Pseudobdellovibrionaceae*.

**Figure 2.**
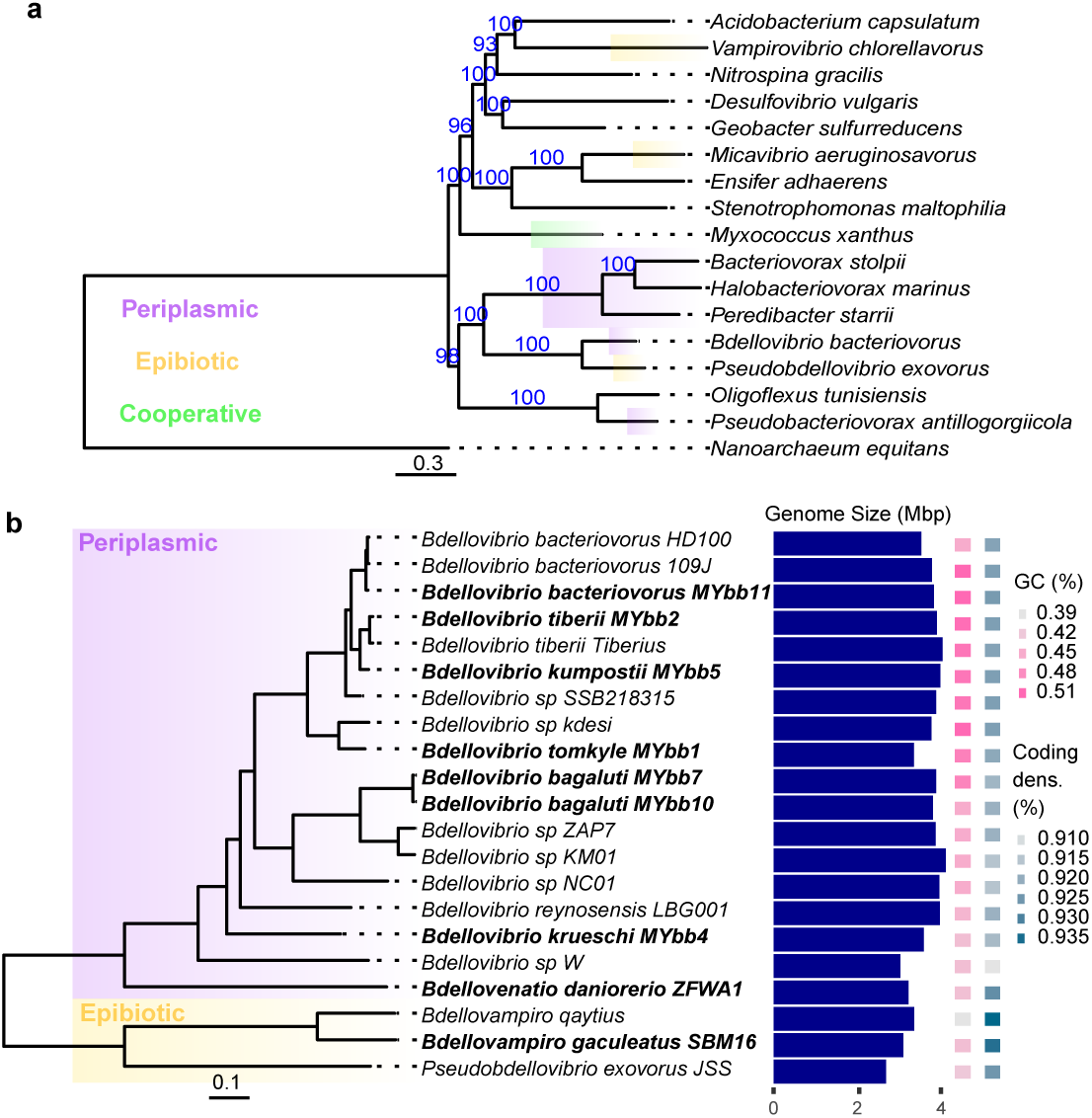
Phylogenetic relationship between various predatory and non-predatory prokaryotes. IQ-TREE was used to infer phylogenetic trees by maximum likelihood with bootstrap approximation. a) Phylogenetic relationships between selected prokaryotic species based on phylogenomic analysis using 265 single-copy genes found in 82.4% of species predicted by Orthofinder. All branch support values are reported in blue. b) Relationships among *Pseudobdellovibrionaceae* strains based on phylogenomic analysis using 1099 single-copy core genes predicted by Orthofinder. All branch support values from bootstrapping approximation were 100. Novel isolates are bolded and listed by the newly proposed names based on 65% AAI genus and 95% ANI species designations.

We used core-genome sequence phylogenetic analysis to examine the evolutionary relationships between 21 *Pseudobdellovibrionaceae* strains including our nine new isolates. The maximum-likelihood phylogenetic tree was constructed based on the amino acid sequences of 1099 single-copy core genes (Figure 2b). *Pseudobdellovibrionaceae* strains fall into two distinct clades that differentiate based on predation strategy. The novel epibiotic predator isolated from the stickleback fish aquaculture water, here named *Bdellovampiro gaculeatus* strain SBM16, clusters with known epibiotic predators *B. qaytius* and *P. exovorus*. The novel isolate from zebrafish aquaculture water (*Bdellovenatio daniorerio* strain ZFWA1) was the most diverged periplasmic *Pseudobdellovibrionaceae strain*.

### Epibiotic *Pseudobdellovibrionaceae* have distinct characteristics indicative of genome reduction

The predators included in Figure 2a were further explored by comparing genome characteristics (Figure 3). Epibiotic *Pseudobdellovibrionaceae* genomes are smaller, have lower GC content, and exhibit a higher coding density than most periplasmic *Pseudobdellovibrionaceae* genomes (Figure 3a). Within the family, *P. exovorus* has the smallest genome, while *B. qaytius* has the lowest GC content and highest coding density. Additionally, all periplasmic predator genomes have less than 350 COGs categorized as M or I functional categories (Figure 3b), indicative of a loss in genes associated with the production of fatty acids, phospholipids, and peptidoglycan [28]. Within this group, epibiotic predators have even lower numbers of M and I COGs and some of the smallest genomes. With high GC content and a larger genome, the cooperative predator *Myxococcus xanthus* differs from other predatory bacteria (Figure 3).

**Figure 3.**
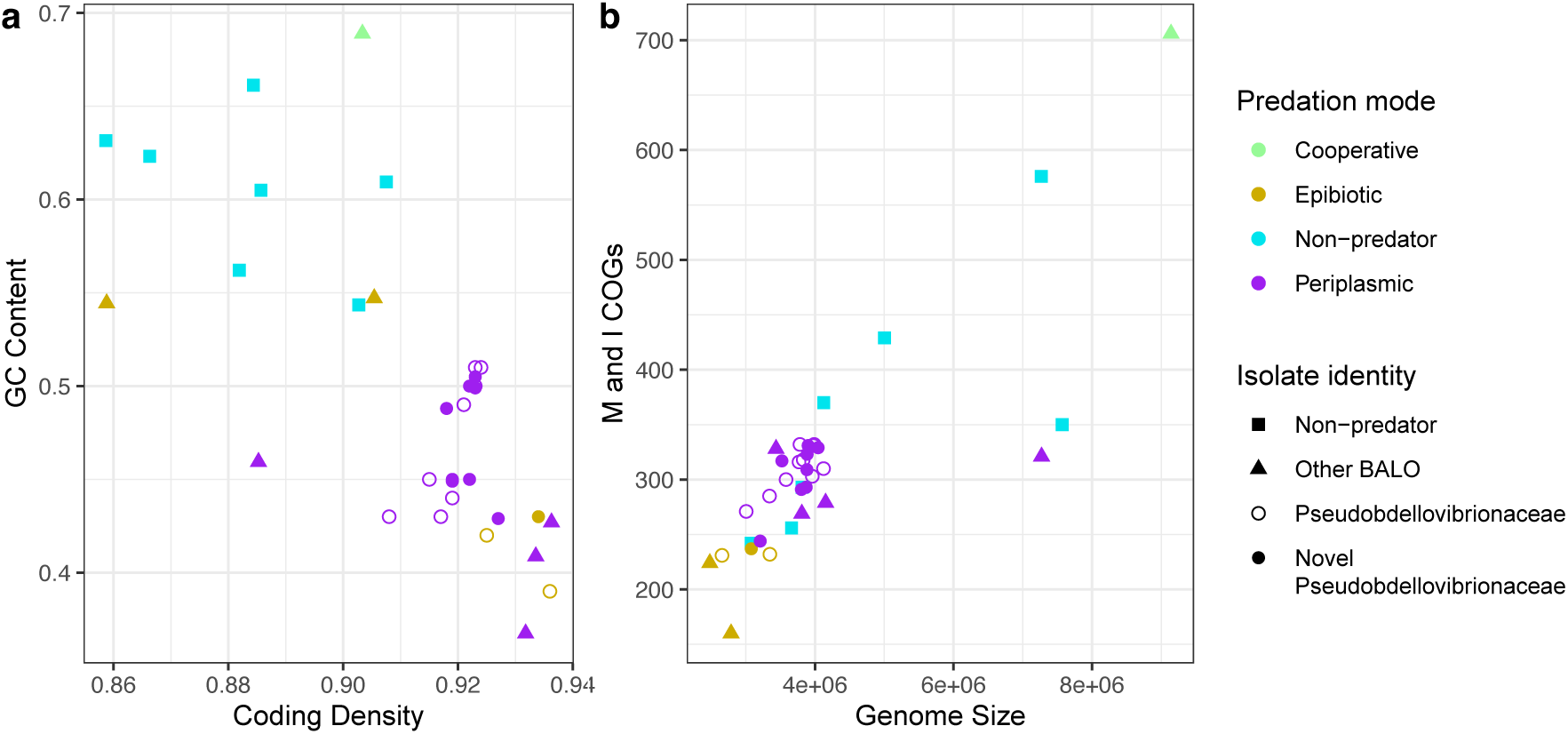
Relationship between different genome metrics of predatory and non-predatory bacteria. a) Relationship between coding density and GC content. b) Relationship between the total combined number of genes in the M (cell envelope biogenesis, outer membrane) and I (lipid metabolism) clusters of orthologous groups (COG) categories and genome size. All bacterial species in Figure 2b are included.

### Periplasmic and epibiotic *Pseudobdellovibrionaceae* vary in gene content

In the comparative genomic analysis of all 21 *Pseudobdellovibrionaceae* strains, between 2616 and 3754 genes were annotated per strain, and 95.7% of genes were identified as belonging to 5897 orthogroups (Table S4, Figure 4a). Genome characteristics of novel strains and type strains are shown in Table 2. A total of 142 of these orthogroups (made up of 354 genes) were exclusive to a single strain. A total of 1360 core orthogroups were found in all genomes.

**Figure 4.**
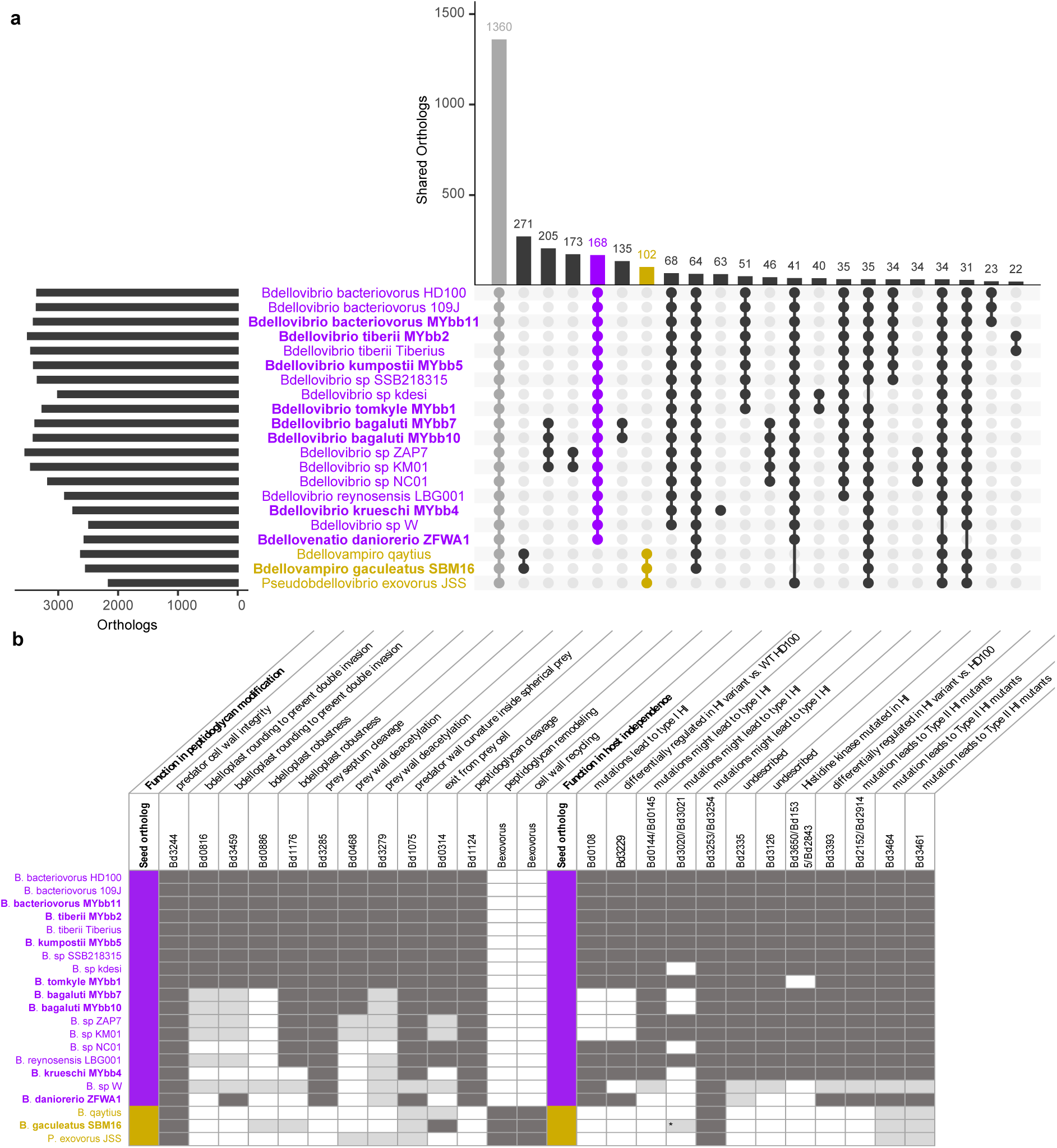
Variation in gene content by predation strategy. a) The number of shared orthologous genes in *Pseudobdellovibrionaceae* strains identified using Orthofinder. The top 22 shared orthologous gene sets are shown. b) Presence of genes involved in peptidoglycan modification and host independence in *Pseudobdellovibrionaceae* isolates. Dark gray cells indicate the presence of the gene with the best protein match listed under seed ortholog. Light gray cells indicate genes with a different seed ortholog that belong in the same orthogroup. The asterisk indicates that only gene Bd3021 is present in SBM16, not Bd3020. All novel isolate names are bolded.

**Table 2.** Genome comparison of novel *Pseudobdellovibrionaceae* isolates with *Bdellovibrio* and *Pseudobdellovibrio* type strains. Genome quality estimates of % completion (% comp.) and % contamination (% cont.) were obtained from Checkm2. The number of coding genes (CDS) and different types of RNA genes were predicted using Prokka. Exclusively present orthogroups (OG) defined by Orthofinder were only found in a single genome, and exclusively absent orthogroups were found in all genomes but one. Novel isolates are bolded, and type strains are indicated with an asterisk.

|  | Genome size | GC content (%) | # of contigs | % comp. | % cont. | CDS | tRNA | rRNA | Other RNA | Exclusively present OG | Exclusively absent OG |
| --- | --- | --- | --- | --- | --- | --- | --- | --- | --- | --- | --- |
| *Bdellovibrio bacteriovorus* HD100* | 3,782,950 | 50.6 | 1 | 99.99 | 0.15 | 3,563 | 36 | 6 | 1 | 0 | 1 |
| ***Bdellovibrio bacteriovorus* strain MYbb11** | 3,901,330 | 50.5 | 1 | 100 | 0.34 | 3,647 | 36 | 6 | 1 | 0 | 0 |
| ***Bdellovibrio tiberii* strain MYbb2\*** | 4,045,310 | 49.8 | 19 | 100 | 0.34 | 3,821 | 35 | 3 | 1 | 3 | 1 |
| ***Bdellovibrio kumpostii* strain MYbb5\*** | 3,886,825 | 49.9 | 1 | 100 | 0.28 | 3,672 | 35 | 6 | 1 | 1 | 0 |
| ***Bdellovibrio tomkyle* strain MYbb1\*** | 3,884,186 | 48.8 | 1 | 100 | 0.25 | 3,702 | 37 | 6 | 1 | 2 | 1 |
| ***Bdellovibrio bagaluti* strain MYbb7\*** | 3,803,878 | 45 | 1 | 99.99 | 0.88 | 3,689 | 34 | 6 | 1 | 1 | 0 |
| ***Bdellovibrio bagaluti* strain MYbb10** | 3,873,366 | 44.9 | 1 | 99.99 | 0.84 | 3,754 | 34 | 6 | 1 | 0 | 0 |
| ***Bdellovibrio krueschi* strain MYbb4\*** | 3,522,424 | 45.2 | 5 | 99.95 | 11.38<sup>a</sup> | 3,479 | 38 | 6 | 2 | 63 | 7 |
| ***Bdellovibrio daniorerio* strain ZFWA1\*** | 3,209,714 | 42.9 | 4 | 99.98 | 0.79 | 3,082 | 38 | 6 | 1 | 17 | 31 |
| ***Bdellovampiro gaculeatus* strain SBM16\*** | 3,080,354 | 43.5 | 16 | 97.12 | 0.76 | 2,885 | 31 | 4 | 1 | 3 | 2 |
| *Pseudobdellovibrio exovorus* JSS* | 2,657,893 | 41.9 | 1 | 99.99 | 0.4 | 2,616 | 33 | 3 | 0 | 14 | 64 |

<sup>a</sup>Other programs reported deviating levels of contamination, including checkm1 (5.41%), BUSCO (3.2%), and NCBI fcs-gx (0%)

Excluding these core orthogroups, *B. qaytius* and *B. gaculeatus* strain SBM16 had the greatest number of unique, shared orthogroups (271). *B. krueschi* strain MYbb4 had the greatest number of strain-specific orthogroups (63) comprising 3.9% of total genes, followed by *B. daniorerio* strain ZFWA1 with 17 strain-specific orthogroups (Table 2). *P. exovorus* JSS had the greatest number of exclusively absent orthogroups (i.e. found in all genomes but *P. exovorus* JSS) (64), followed by *B. daniorerio* strain ZFWA1 with 31 exclusively absent orthogroups.

Epibiotic strains shared 102 unique orthogroups that had a higher relative frequency of genes involved in the transport and metabolism of amino acids, nucleotides, coenzymes, and lipids as well as in secondary metabolite biosynthesis, intracellular trafficking, secretion, and vesicular transport (Figure S3). Orthogroup eggNOG-mapper annotations are listed in Table S5. Epibiotic predators also exclusively encoded some of the genes involved in folate biosynthesis (including *folB*, *folP*, and *folC*), as well as ABC transporters of substrates across the cytoplasmic membrane (including *dppB*), synthesis of glutamate (*glnN*) and purine (*purK*), and disposal of toxic ammonia (*rocF*). Orthogroups unique to periplasmic predators (168) had a higher relative frequency of genes involved in numerous COG categories including transcription, replication, recombination and repair, cell wall biogenesis, cell motility, and signal transduction mechanisms (Figure S3). Here, unique genes included genes encoding histidine kinases and genes involved in chemotaxis.

The distribution of several peptidoglycan-modifying enzymes differs between predation strategies. While all *Pseudobdellovibrionaceae* encode a housekeeping penicillin binding protein (*dacB*) involved in cell wall integrity, periplasmic predators exclusively share an *mltA*-like lytic transglycosylase (Bd3285) and a *slt* lytic transglycosylase (Bd1124) (Figure 4b). Epibiotic strains encode two peptidoglycan modifying enzymes that were absent from periplasmic strains (Figure 4b). These include anhydro-N-acetylmuramic acid kinase (anmK) and a polysaccharide deacetylase involved in cell wall recycling and peptidoglycan remodeling, respectively.

### Many *Pseudobdellovibrionaceae* lack genes involved in host independence

Genomes of the epibiotic predators and *Bdellovibrio bagaluti* strains MYbb7 and MYbb10, as well as *Bdellovibrio* sp. ZAP7 and KM01, lack the *hit* locus gene Bd0108 (Figure 4b). A similar pattern was observed for the LysR family transcriptional regulator gene Bd3229. However, this gene was also absent in *B. bacteriovorus* W.

Further, epibiotic predators and *Bdellovenatio daniorerio* strain ZFWA1 lack the genes *motAB1* (Bd0144, Bd0145) and *motAB2* (Bd3021, Bd3020 except for SBM16), and a histidine kinase (Bd3126), which are genes previously reported to be involved in host independence (HI; i.e., prey-independent feeding, see discussion). In contrast, two other putative HI genes, *rhlB* (Bd3461) and *pcnB* (Bd3464), as well as *motAB3* (Bd3254 and Bd3253) were present in all strains. Additionally, the described HI-related histidine kinase Bd3650/Bd1535/Bd2843 was absent in all these strains, and also in MYbb1 and *B. bacteriovorus* W. Two other histidine kinase genes (Bd3393 and Bd2335) and Bd2152 (a glycerol-3-phosphate transporter [29]) were exclusively absent in epibiotic predators. Additionally, epibiotic predators and ZFWA1 exclusively share 12 orthologous genes with annotated functions involved in initial binding of peptides in periplasmic space (*dppA*) for ABC-type transport, methionine salvage (*mtnA* and *mtnP*), cell wall formation (*murB*), and tyrosine metabolism (*hgo*).

### *Pseudobdellovibrionaceae* strains show indications of habitat specificity

We found ASVs matching the 16S rDNA of our newly isolated strains in host-microbiome datasets of zebrafish, stickleback, and worms (Table 3). Interestingly, zebrafish microbiomes only contained *Pseudobdellovibrionaceae* ASVs matching strains that had been isolated from aquatic habitats. In contrast, stickleback microbiomes contained a more diverse set of *Pseudobdellovibrionaceae*, including strains that had been isolated from the direct environment of zebrafish (ZFW), stickleback (SBM), and worms (MYbbs), and were present in up to 80% of samples. Lastly, worm microbiomes rarely contained *Pseudobdellovibrionaceae* ASVs (3% of samples) and did not harbor *Pseudobdellovibrionaceae* ASVs associated with zebrafish or stickleback. However, they did contain *Pseudobdellovibrionaceae* ASVs that had been previously associated with aquatic habitats, although most of the *Pseudobdellovibrionaceae* ASVs belonged to strains isolated from worm-associated environments or other terrestrial habitats.

**Table 3:**
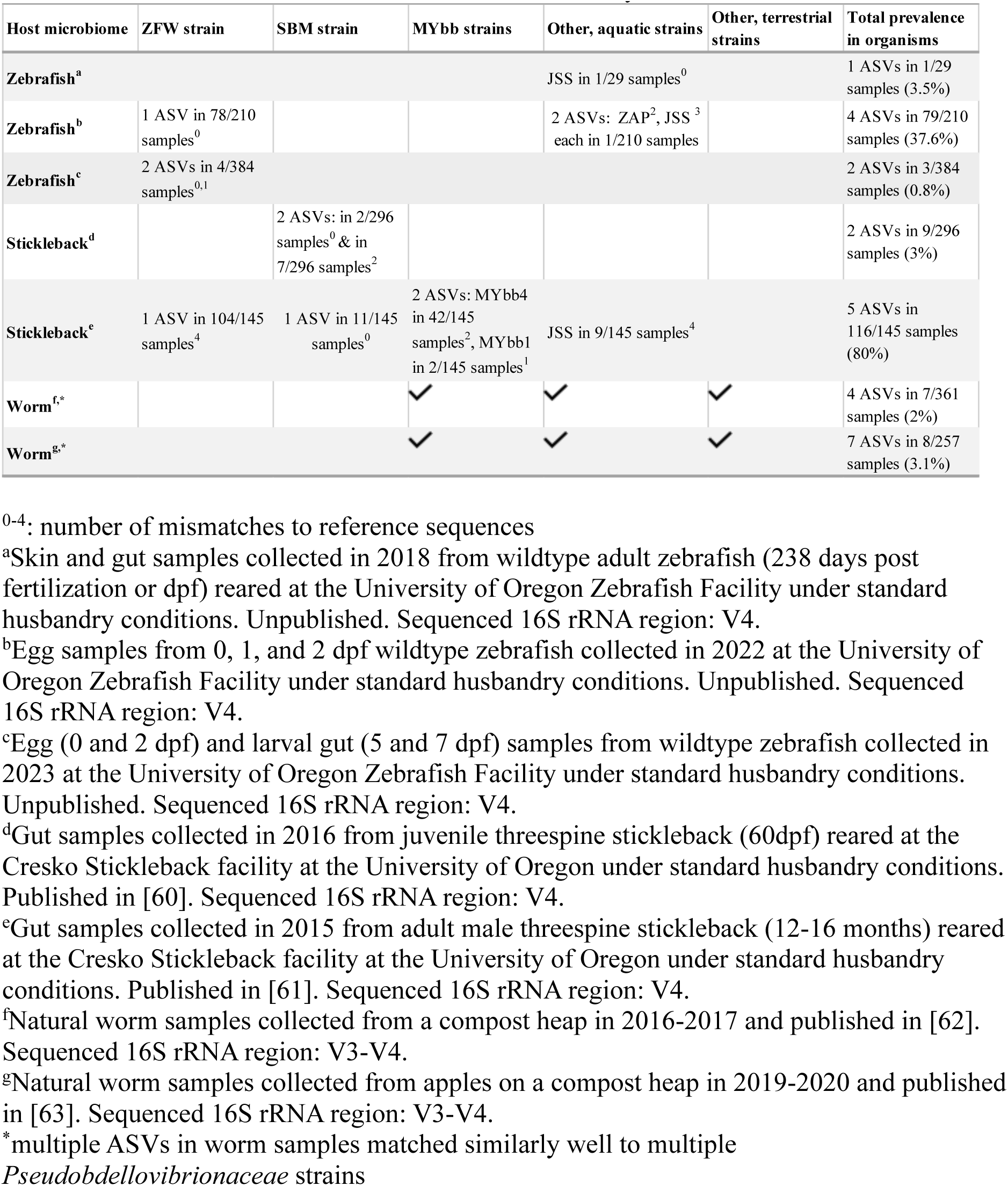
*Pseudobdellovibrionaceae* strains can be found in 16S rDNA data sets of relevant hosts. Number of the different *Pseudobdellovibrionaceae* ASVs with up to 4 mismatches (i.e. 99% identity) and their prevalence are shown for each microbiome study and all *Pseudobdellovibrionaceae* strains used in this study.

## Discussion

*Bdellovibrio* and like organisms (BALOs) play an important role in microbial communities [30–32] and there is growing interest in using BALOs to manipulate microbial communities, especially within host organisms [33–35]. To broaden our understanding of BALO diversity, especially those associated with animal hosts, we isolated nine novel *Pseudobdellovibrionaceae* strains from the environments of three important model animals: the zebrafish, threespine stickleback fish, and the nematode *C. elegans*. We characterized these strains using TEM, comparative genomics and phylogenetic analysis. Two of these nine strains were sufficiently diverged genetically to be considered new genera, including only the third *Pseudobdellovibrionaceae* strain known to use an epibiotic predation strategy, while six strains can be grouped into five new species within the genus *Bdellovibrio.* Our phylogenetic analysis indicates that there are two clades within the family *Pseudobdellovibrionaceae*, separating periplasmic from epibiotic genera (Figure 2b). These novel isolates significantly expand our understanding of BALO biology, especially the genetics and evolution of predation strategies.

### Epibiotic predation is a derived trait among BALOs

Phylogenetic analysis of our novel isolates allowed us to better understand the evolution of predation modes among BALO species. Bacterial predation is not a monophyletic trait, and predatory bacteria are found in other bacterial phyla. When including other bacterial phyla in the broader phylogeny (Figure 2a), we identified a single clade that contained all periplasmic BALO genera, suggesting that periplasmic predation may be the ancestral phenotype. In contrast, epibiotic BALO genera are dispersed amongst non-predators and periplasmic predators alike, suggesting that epibiotic predation arose in multiple independent lineages. This evolutionary hypothesis differs from that of Deeg et al. [16], who proposed that epibiotic predation is ancestral and periplasmic predation the derived state within the *Pseudobdellovibrionaceae*.

However, Deeg et al. [16] used a single gene (16S rRNA) to infer their phylogeny, while we used a set of over 250 single-copy core genes.

Our comparative genomic analyses provided additional support for our hypothesis that periplasmic predation is ancestral. Pasternak et al. [14] noted genome reduction in periplasmic predators driven by the dependence on prey amino acids and vitamins as is observed in host-dependent bacteria [36]. If periplasmic predation is ancestral, derived epibiotic predators therefore would exhibit a secondary phase of genome streamlining as first hypothesized by Pasternak et al. [15]. In the present study, epibiotic *Pseudobdellovibrionaceae* genomes show signs of genome degradation, similar to what has been shown for host-associated symbionts [28]. This includes higher AT content and coding density, smaller genome size, including fewer DNA repair genes and fewer genes involved in cell envelope biosynthesis and morphology when compared to non-predators (Figure 3 and Figure S3). However, we can base this interpretation only on observations of the three epibiotic *Pseudobdellovibrionaceae* strains described so far.

However, this streamlining of epibiotic predators seems to be restricted to *Pseudobdellovibrionaceae* species as periplasmic BALOs in the family *Bacteriovoracaceae* including *Halobacteriovorax marinus*, *Bacteriovorax stolpii*, and *Peredibactor starii* had low GC content and high coding density comparable to epibiotic *Pseudobdellovibrionaceae*.

Likewise, epibiotic BALOs *Micavibrio aeruginosavorus* and *Vampirovibrio chlorellavorus* had higher GC content and lower coding density than all *Pseudobdellovibrionaceae* predators (Figure 3b).

### The evolution of epibiotic predation may have involved the loss of “host independence” genes

Our comparative genomic analysis identified multiple genes that may have been lost in epibiotic predators during their evolution from a periplasmic ancestor. Many of these are genes involved in “host independence” in *Bdellovibrio bacteriovorus* (Figure 4b), a prey-independent feeding strategy that may be induced under limited prey conditions in nature and on protein-rich media in the laboratory [6, 37] and is caused by a mutation in any of a number of genes. We identified multiple genes associated with host independence (specifically, but not exclusively Type I host independence; [38–43]), including Bd3126 (a histidine kinase), and one of the three motAB gene clusters, Bd0144/Bd0145 (*motAB1*), that were absent in the epibiotic *Pseudobdellovibrionaceae* species and their closest relative that used periplasmic predation (*Bdellovenatio daniorerio* ZFWA1), but were present in other periplasmic predators (Figure 4b).

### Functional differences in peptidoglycan modification differentiate predation strategies

Our comparative analyses provided novel information regarding the biochemistry underlying different predation strategies and builds upon previous studies [15, 18]. Epibiotic genomes lack some peptidoglycan modifying enzymes that have been shown to be important in periplasmic predation (Figure 4B; [16]). Assuming periplasmic predation as the ancestral state, this likely represents a gene loss for epibiotic predators. All *Pseudobdellovibrionaceae* strains encode *dacB*, a housekeeping PBP4 (penicillin binding protein) likely involved in the maintenance of the predator’s own cell wall [44]. Periplasmic *Bdellovibrio* predators in the clade that contains both strains HD100 and MYbb1 (Table 2a) all possess a suite of enzymes involved in the modification of the prey cell wall. For example, two *dacB*-like genes, Bd0816 and Bd3459, involved in prey cell rounding to form an osmotically stable bdelloplast and reducing the rate of secondary invasions [44], were absent from some more diverged periplasmic strains and all epibiotic strains. The rate of wasteful secondary invasions may be higher in strains that lack these genes, but this is currently unknown. Similarly, L,D-transpeptidases, Bd0886 and Bd1176, strengthen the prey bdelloplast wall to resist bursting during predator growth within the prey cell [45]. While absent from various strains, an ortholog of these genes is encoded in *B. gaculeatus* strain SBM16. Since bdelloplast formation is not necessary for epibiotic predators, these genes may perform a different role in SBM16 or may be fated for removal in further genome streamlining. Interestingly, all four genes involved in bdelloplast integrity were absent from *B. krueschi* strain MYbb4. This, together with the abundance of exclusively present orthogroups (Table 2) may reflect MYbb4’s evolved specificity to predate *Ochrobactrum*, although host range was not exhaustively examined here. Further, the diversity, distribution, and function of predatory peptidoglycan-modifying genes is likely influenced by structural and biochemical properties of the prey cell wall such as cross-link chemistry [45], bacterial capsules, surface layers, or acetylation state [46].

Notably, an *mltA*-like lytic transglycosylase is exclusive to periplasmic predators. In *Bdellovibrio bacteriovorus* strain HD100, this protein is secreted into the prey periplasm and cleaves the peptidoglycan septum of dividing prey, facilitating the conversion of actively dividing prey into a single spherical bdelloplast [47]. All strains contain an LD carboxypeptidase involved in predator cell wall curvature which causes the formation of a vibrioid and facilitates efficient intracellular growth of HD100 in a spherical prey niche [48]. What role this plays in epibiotic strains is yet to be determined. Most periplasmic predators and *P. exovorus* strain JSS encode peptidoglycan deacetylases (Bd0468 and Bd 3279 or orthologs) that deacetylate the peptidoglycan of the prey [49]. This allows the predator to differentiate between the cell wall of the prey and that of the predator itself with the enzyme dslA (Bd0314) which cuts peptidoglycan depending on acetylation state and functions as an exit enzyme by destroying the prey cell wall [50]. Strains MYbb4, ZFWA1, and JSS all lack *dslA*, suggesting that MYbb4 and ZFWA1 possess an alternative exit strategy.

Epibiotic *Pseudobdellovibrionaceae* strains exclusively share *anmK*, which encodes a bifunctional glycosidase/kinase involved in recycling of cell-wall derived anhydroMurNAc [51]. Interestingly, all non-*Pseudobdellovibrionaceae* epibiotic predators also encode AnmK as do non-predator relatives (*A. capsulatum*, *V. chlorellavorus*, *M. aeruginosavorus*, *E. adhaerens*, and *S. maltophilia*). While not present in periplasmic predators, this widely conserved gene may be integral to epibiotic predation. Cell-wall recycling through MurNAc dissimilation is a pathway by which bacteria can utilize peptidoglycan fragments from the environment or from the endogenous cell wall [51]. This could be advantageous for predatory bacteria to scavenge prey cell wall material [52]. However, *murQ* is also necessary for anhydroMurNAc recycling, but is absent in *Pseudobdellovibrionaceae* strains. Both *anmK* and *murQ* are present in epibiotic predators *M. aeruginosavorus* and *V. chlorellavorus* as well as *E. adhaerens*. While some epibiotic predators may be able to utilize prey cell wall material through this process, the role and exclusivity of *anmK* in epibiotic *Pseudobdellovibrionaceae* is yet to be uncovered.

### The family *Pseudobdellovibrionaceae* is more genus and species rich than previously thought

Using ANI, we identified new *Pseudobdellovibrionaceae* strains belonging to five new species and two new genera. Further, existing strains proposed to belong to *B. bacteriovorus* likely represent separate species. Renaming these strains as distinct species would enhance our understanding of *Pseudobdellovibrionaceae* diversity and evolution.

Targeting few environments in relation to host organisms of interest was enough to isolate novel BALOs. Thus, true BALO species diversity is largely underexplored. This may be particularly true for difficult to culture species such as epibiotic predators. In general, epibiotic predators have a narrower prey range than periplasmic predators and produce fewer offspring per prey cell, resulting in lower population growth rates [13, 16, 17], a pattern we also observed for our novel epibiotic isolate B. gaculeatus strain SBM16.

### *Pseudobdellovibrionaceae* are likely widespread among host microbiomes

Although there is growing interest in using BALOs to manipulate host-associated microbiomes, there have been very few studies that have attempted to isolate host-associated BALO species [53–57]. By sampling the immediate environment of three model animal hosts (zebrafish, threespine stickleback, and *C. elegans*) we successfully isolated new *Pseudobdellovibrionaceae* and detected ASVs (highly) similar to the 16S rDNA sequences of those strains in microbiomes of the hosts: *Bdellovenatio daniorerio* strain ZFWA1 ASVs were found in zebrafish eggs and larvae, as well as in samples of threespine stickleback guts (Table 3). Zebrafish microbiomes exclusively contained *Pseudobdellovibrionaceae* ASVs of strains originally isolated from aquatic environments. In contrast, stickleback microbiomes contained *Pseudobdellovibrionaceae* ASVs of species originally isolated from many different environments (Table 3), including two *Bdellovibrio* species that we isolated from compost (*B. tomkyle* strain MYbb1 and *B. krueschi* strain MYbb4). Interestingly, ASVs of *B. krueschi* strain MYbb4 were highly prevalent across stickleback gut samples. MYbb4 specifically preys on *Ochrobactrum*, and we indeed detected *Ochrobactrum* ASVs in these same stickleback samples. We found ASVs of aquatic *Pseudobdellovibrionaceae* in worm microbiomes (Table 3). One example is the Tiberius strain, isolated from the nutrient-rich Tiber River in Italy. This strain belongs to the same species as *B. tiberii* MYbb2, which we isolated from compost material in Kiel, Germany. This indicates that the same *Bdellovibrio* strain can be found in multiple habitats and suggests that habitat preferences may depend more on the presence of suitable prey than on the habitat itself.

The prey preferences of different *Pseudobdellovibrionaceae* species likely influences their distributions. For example, *B. daniorerio* strain ZFWA1 was isolated using *Aeromonas veronii*, and this genus is commonly found associated with fish as well as in free-living aquatic microbiomes. We observed ASVs of *B. daniorerio* in zebrafish and stickleback microbiomes, which also contained *Aeromonas* species. Similarly, most of our compost (MYbb) isolates were isolated using *E. coli* as prey. *E. coli* strains are widespread across different environments, and indeed we find many *Pseudobdellovibrionaceae* ASVs in microbiomes from both worms and stickleback fish. In contrast, ASVs of *B. gaculeatus* strain SBM16 were exclusively present in stickleback microbiomes. This species was isolated from a stickleback aquaculture facility, a moderately saline environment. We observed that this species had higher population growth rates in the laboratory when a saline medium was used. However, further research is needed to confirm whether this strain truly exhibits strong habitat specificity toward saltwater.

Interestingly, this strain had a distinct flagellum morphology compared to the other isolates (i.e., smaller wave amplitudes), which may have functional implications in saline environments. In *Shewanella*, a smaller wave amplitude led to decreased spreading in soft agar (consistent with our observation that SBM16 does not grow well on double-layer agar plates) but increased velocity at higher viscosity (as in salt water) [58], a phenomenon observed in other bacteria as well [59].

Isolating several new *Pseudobdellovibrionaceae* strains, including an epibiotic BALO, revealed that the diversity within the family is highly underestimated and helped improve genome-based predictions regarding predation mode and its evolutionary origin. Efforts in isolating and sequencing new predatory bacteria will further enhance our knowledge and allow for the testing and application of these strains as alternative probiotics or antibiotics in the future.

Formal description of five new species and two new genera within the *Pseudobdellovibrionaceae*

*Bdellovibrio tiberii* sp. nov. (/tiˈbeːri.iː/, derived from the name of the river Tiber, from which the first strain was isolated) with type strain MYbb2^T^

Highly motile cells of vibrioid shape (0.25 ± 0.04 µm^2^ in size) with a single flagellum (3.34 ± 0.36 µm in length) and a periplasmic predation strategy. The type strain was isolated from *C. elegans*-associated compost material, collected from the Kiel Botanical Garden using *Escherichia coli* ML35 as prey. Cells grow on prey lawns as plaques at 28°C.

The species belongs to the genus *Bdellovibrio* (within the family *Pseudobdellovibrionaceae*). The genome has a G+C content of 49.8% and is approximately 4.05 Mb in size. The strain is accessible at the DSMZ () and the ATCC ().

*Bdellovibrio kumpostii* sp. nov. (/kumˈpɔs.ti.iː/, latinized from the German word Kompost describing the isolation source, a compost heap in Kiel, Germany) with type strain MYbb5^T^ Highly motile cells of vibrioid shape (0.21 ± 0.03 µm^2^ in size) with a single flagellum (3.02 ± 0.55 µm in length) and a periplasmic predation strategy. The type strain was isolated from *C. elegans*-associated compost material, collected from the Kiel Botanical Garden using *Escherichia coli* ML35 as prey. Cells grow on prey lawns as plaques at 28°C.

The species belongs to the genus *Bdellovibrio* (within the family *Pseudobdellovibrionaceae*). The genome has a G+C content of 49.9% and is approximately 3.89 Mb in size. The strain is accessible at the DSMZ () and the ATCC ().

*Bdellovibrio tomkyle* sp. nov. (/tɔmˈki.le/, derived from the Low German binomial tom kyle referring to the old name for the city of Kiel in Germany, meaning tom “at the” kyle “fjord”, as the strain was isolated in Kiel) with type strain MYbb1^T^

Highly motile cells of vibrioid shape (0.19 ± 0.02 µm^2^ in size) with a single flagellum (3.02 ± 0.22 µm in length) and a periplasmic predation strategy. The type strain was isolated from *C. elegans*-associated compost material, collected from the Kiel Botanical Garden using *Escherichia coli* ML35 as prey. Cells grow on prey lawns as plaques at 28°C.

The species belongs to the genus *Bdellovibrio* (within the family *Pseudobdellovibrionaceae*). The genome has a G+C content of 48.8% and is approximately 3.88 Mb in size. The strain is accessible at the DSMZ () and the ATCC ().

*Bdellovibrio bagaluti* sp. nov. (/baɡaˈluːti/, derived from the Low German word Bagalut, referring to someone who is mischievous or unruly to reflect the predatory nature of the strain) with type strain MYbb7^T^

Highly motile cells of vibrioid shape (0.21 ± 0.03 µm2 in size) with a single flagellum (2.99 ± 0.44 µm in length) and a periplasmic predation strategy. The type strain was isolated from *C. elegans*-associated compost material, collected from the Kiel Botanical Garden using *Escherichia coli* ML35 as prey. Cells grow on prey lawns as plaques at 28°C.

The species belongs to the genus *Bdellovibrio* (within the family *Pseudobdellovibrionaceae*). The genome has a G+C content of 45% and is approximately 3.8 Mb in size. The strain is accessible at the DSMZ () and the ATCC ().

*Bdellovibrio krueschi* sp. nov. (/ˈkʁʏʃi/, derived from the Low German word krüsch which describes someone who is picky reflecting the small prey range of the species) with type strain MYbb4^T^

Highly motile cells of vibrioid shape (0.25 ± 0.03 µm2 in size) with a single flagellum (2.58 ± 0.33 µm in length) and a periplasmic predation strategy. The type strain was isolated from *C. elegans*-associated compost material, collected from the Kiel Botanical Garden using *Ochrobactrum* MYb71 as prey. Cells grow on prey lawns as plaques at 28°C. So far, only members of the genus *Ochrobactrum* have been identified as prey.

The species belongs to the genus *Bdellovibrio* (within the family *Pseudobdellovibrionaceae*). The genome has a G+C content of 45.2% and is approximately 3.52 Mb in size. The strain is accessible at the DSMZ () and the ATCC ().

*Bdellovenatio* gen nov. (/bdɛloʊvəˈneɪʃiˌoʊ/, derived from Latin *bdella* meaning “leech” and *venatio* meaning “hunting”, referring to the rapid movement of the cells) with type species *daniorerio* sp. nov. (/dæniɔːˈrɛrioʊ/, derived from the Latin binomial *Danio rerio,* the scientific name of the zebrafish in which this bacterium has been found) and type strain ZFWA1^T^

Highly motile cells of vibrioid shape (0.25 ± 0.05 µm^2^) with a single flagellum (2.77 ± 0.25 µm in length) and a periplasmic predation strategy. The type strain was isolated from water collected at the University of Oregon zebrafish aquaculture facility on the campus in Eugene, Oregon, USA, using *Aeromonas veronii* ZOR0001as prey. Cells grow on prey lawns as plaques at 28°C. The species belongs to a unique genus in the family *Pseudobdellovibrionaceae*. The genome of has a G+C content of 42.9% and is approximately 3.21 Mb in size. The strain is accessible at the DSMZ () and the ATCC ().

*Bdellovampiro* gen nov. (/bdɛloʊˈvæmpɪroʊ/, derived from Latin *bdella* meaning “leech” and *vampiro* meaning “vampire” referring to its epibiotic predation strategy) with type species *gaculeatus* sp. nov. (/gɑːkjuːˈliːətəs/, a contraction of the Latin binomial *Gasterosteus aculeatus,* the scientific name of the threespine stickleback fish in which this bacterium has been found) and type strain SBM16^T^

Highly motile cells of elongated vibrioid shape (0.77 ± 0.24 µm^2^) with a thin and very long (7.13 ± 1.57 µm) single flagellum and an epibiotic predation strategy. The type strain was isolated from water collected at the University of Oregon threespine stickleback aquaculture facility on the campus in Eugene, Oregon, USA, using *Escherichia coli* ML35 as prey. Cells grow together with prey at 20°C. This strain is mesohalophilic, growing optimally in media amended with 2 g/L sea salt. The species belongs to a unique genus in the family *Pseudobdellovibrionaceae*. The genome has a G+C content of 43.5% and is approximately 3.08 Mb in size. The strain is accessible at the DSMZ () and the ATCC ().

## Acknowledgements

We thank the Bohannan and Schulenburg groups for discussions and advice on this work. For genomic sequencing, we further thank the Competence Centre for Genomic Analysis (CCGA) Kiel. For obtaining environmental samples, we thank the Botanical Garden in Kiel, Germany, the Cresko Stickleback Facility, and the University of Oregon Zebrafish Facility. We are grateful for funding from the German Science Foundation (Deutsche Forschungsgemeinschaft, DFG) within the Collaborative Research Center (CRC) 1182 on the Origin and Function of Metaorganisms (Project-ID 261376515 – SFB 1182, project A4.3 to HS), individual DFG project JO 1786/1-1 (to JJ), the Max-Planck Society (fellowship to HS), the NIH P01-GM125576 to BJMB, and the NIH Research Supplement to Promote Diversity in Health-Related Research to RLM.

## Data Availability Statement

All raw sequences are available in the SRA under PRJNA1185762 and assembled *Pseudobdellovibrionaceae* genomes are available in the NCBI under PRJNA1185297.

## Supplementary Tables

**Table S1. Size measurements for *Pseudobdellovibrionaceae* strains described in this study.** Measurements of bacteria were performed on images taken at 11,000x (TIA camera) or 16,500x (MegaView III camera) magnification using FIJI software (Schindelin et al., 2012). A segmented line tool was used to measure the length of flagella, and a polygon tool was used to measure the cell body area, perimeter and Feret diameter. Culture column indicates whether fresh prey was added before imaging or whether standard overnight cultures were used. The mean, standard deviation (SD) and count numbers are displayed for flagellum and cell body measurements. Units for each measurement are indicated in square brackets. Count column refers to the number of images analyzed for mean values. Run column indicates how many independent experiments (the first number) and how many independent grids (the second number) were analyzed for each sample.

**Table S2. Average Nucleotide Identity (a) and Average Amino Acid Identity (b) between strains.** Pairwise ANI was calculated with fastANI v1.34 and pairwise AAI was calculated with CompareM v0.1.2.

**Table S3. Genbank accession numbers of genome assemblies included in phylogenomic analyses.** Prokaryotic genomes were used to infer phylogenetic relationships between periplasmic and epibiotic BALOs. *Pseudobdellovibrionaceae* genomes were used to infer phylogenetic relationships between periplasmic and epibiotic existing and novel *Pseudobdellovibrionaceae* strains.

**Table S4. Orthofinder output of gene counts by *Pseudobdellovibrionaceae* isolate in each orthogroup.** Rows are predicted orthogroups and columns are genome assemblies. Counts represent the number of gene calls from a genome that belong to a specific orthogroup.

**Table S5. EggNOG-mapper annotations of each Prodigal-predicted gene in Orthofinder- predicted orthogroups.** Each row represents a unique gene identified by Query. Each gene is annotated from a specific *Pseudobdellovibrionaceae* genome and belongs to an orthogroup. Orthogroups may have multiple genes from the same and/or different genomes. Each query has a Seed ortholog that is the best matching sequence in the eggNOG protein space. Genome id is the Prodigal-assigned id to the genome assembly. Input into eggNOG-mapper was the Prodigal faa file output of each genome assembly. Genome names reflect current database names.

## Supplementary Figures

**Figure S1: Cell shape and predation mode of periplasmic *Pseudobdellovibrionaceae* strains MYbb1, MYbb5, MYbb7, MYbb10 and MYbb11.** Stages in the life cycle of periplasmic *Pseudobdellovibrionaceae* isolates are shown on top. Numbers in image corners correspond to the proposed life cycle stage. Stage 1 images represent *Pseudobdellovibrionaceae* cells in attack phase. In stage 2, predators are attached to prey cells (p). After penetration of the prey’s outer membrane, *Pseudobdellovibrionaceae* cells enter stages 4-7 with intraperiplasmic growth in bdelloplasts (bd), with one or more predator cells confined within a prey cell (p). Samples were imaged by TEM after negative staining. Scale bar is 500 nm. Life cycle images created in BioRender. Wülbern, J. (2024) BioRender.com/t31h353.

**Figure S2. Average Nucleotide Identity (a) and Average Amino Acid Identity (b) between strains.** Pairwise ANI was calculated with fastANI v1.34 and pairwise AAI was calculated with CompareM v0.1.2. Both were visualized in R with ComplexHeatmap v2.20.0. Novel isolates are listed in bold.

**Figure S3. Relative frequency of COG categories in orthogroups unique to the core, epibiotic, and periplasmic genomes.** Orthogroups were annotated with a COG category using eggNOG-mapper and classified as core (found in all genomes), epibiotic, or periplasmic. Total orthogroups per classification are listed in the figure legend. Relative frequency is calculated by summing COG annotations per category within a classification and divided by the total number of orthogroups per classification.

