## Supplementary Methods, Figures and Tables for "Comparative analysis of novel *Pseudobdellovibrionaceae* genera and species yields insights into the genomics and evolution of bacterial predation mode"

*Isolation and cultivation of new* Pseudobdellovibrionaceae *strains*

The novel strains we describe were isolated from the laboratory husbandry environments associated with zebrafish and threespine stickleback, and the natural environment of *Caenorhabditis elegans* (compost, i.e. degraded plant material).

a) *Fish aquaculture derived* Pseudobdellovibrionaceae *isolation*

*Pseudobdellovibrionaceae* strains were enriched from water collected from fish aquaculture facilities at the University of Oregon in Eugene, Oregon, USA. Water was collected from the sump tank of the Cresko Stickleback facility in November 2021 and sentinel tanks of the University of Oregon Zebrafish Facility in April 2022. To remove debris and larger organisms, the water samples were initially filtered through 0.45μm filters. Subsamples of filtered water (20-100ml) were transferred to sterile flasks and incubated for several days at 28°C and 180rpm. To enrich for BALOs, water samples were inoculated with potential prey, either *E. coli* ML35 (for the stickleback facility water) or *Aeromonas veronii* ZOR0001 (for the zebrafish facility water). ZOR0001is a strain of *Aeromonas veronii* isolated from the zebrafish gut [1, 2]. Prior to inoculation into the BALO enrichments, overnight cultures of prey species were resuspended in HM buffer (25 mM HEPES (N-[2-hydroxyethyl]piperazine-N'-[2-ethanesulfonic acid]) at pH 7.4, 3 mM $CaCl_2 \cdot 2H_2O$ and 2 mM $MgCl_2 \cdot 7H_2O$) [3]. Cultures were visually monitored daily for clearance of prey and via microscopy for the presence of small, highly motile cells (indicative of the presence of BALOs).

To isolate single BALO strains, the cultures were serially diluted in HM buffer and used in plaque assays with the prey strains used for enrichment. Plaque assays were performed as described previously [3], using DNB soft agar (7 g agar/L) and DNB agar plates, along with 200 μl of overnight prey culture resuspended in HM buffer and 100 μl of the BALO enrichment. Top agar plates were incubated at 28°C and checked daily for the presence of plaques. Plaques were aspirated from the top agar with a 200µl pipette, then transferred to 200µl of DNB. After incubating at 28°C for 30 min, the BALO isolates were inoculated with 50µl of prey cells resuspended in HM buffer. These co-cultures were incubated at 28°C and 180rpm. After several days, the presence of small, highly motile cells was confirmed with microscopy. Glycerol stocks (25% (v/v)) were generated and used for subsequent DNA isolation and whole genome sequencing.

### *b) Worm substrate derived* Pseudobdellovibrionaceae *isolation*

All nematode-derived *Pseudobdellovibrionaceae* (MYbb) were isolated from four compost samples associated with *C. elegans*. Compost material was sampled in November 2016 from the Botanical Garden in Kiel, Germany (54.34826809054093 N, 10.116846904750298 E). To isolate BALOs from the compost material, we followed the instructions of Jurkevitch et al. [3], using basic protocol 1 for soil samples and alternate protocol 3 for BALO enrichment. Specifically, an approximately 10-20 g compost sample was incubated in 50 ml sterile water under shaking conditions (200 rpm) at 28°C for multiple days.

To enrich for BALOs that prey on specific microbes, cultures were spiked with either *E. coli* ML35 or known *C. elegans* microbiome members (*Stenotrophomonas* MYb238, *Pseudomonas* MYb187, and *Ochrobactrum* MYb71) suspended in 25 mM N-[2-hydroxyethyl]piperazine-N'-[2-ethanesulfonic acid] supplemented with 3 mM CaCl2·2H2O and 2 mM MgCl2·7H2O (25 mM HEPES, pH 7.4). Clearance of cultures indicated predatory activity. A subsample was subsequently investigated under the microscope to look for small and fast-moving cells.

To isolate single *Pseudobdellovibrionaceae* strains, these cultures were concentrated by centrifugation. First, they were centrifuged at 500 x g for 5 minutes to remove large particles. Second, the supernatant was centrifuged for 20 minutes at 16,000 x g. The supernatant was discarded, and the pellet was resuspended in 5 ml of 1:10 diluted nutrient broth supplemented with 3 mM $CaCl_2 \cdot 2H_2O$ and 2 mM $MgCl_2 \cdot 7H_2O$ (DNB). This suspension was then filtered through a 0.45 µm filter to remove larger bacteria.

The suspension was then serially diluted and used in plaque assays with the same prey bacteria used for spiking. Plaque assays were performed as described previously [3], using DNB soft agar (7 g agar/L) and DNB agar plates, along with 300 µl of prey bacteria (washed with 25 mM HEPES) and 100 µl of BALO suspension. Plates were incubated at 28°C and checked repeatedly over the next few days. Plaques that appeared after 24 hours were considered phage-derived and labeled accordingly. Plaques that appeared later were considered BALO-derived and were subsequently transferred from the agar via a cut 1 ml pipette tip (to increase the diameter of the opening) into an Eppendorf tube containing 500 µl of 25 mM HEPES buffer. These plaque cultures were incubated for 10 minutes at room temperature, vortexed, and examined for small, fast-swimming cells under the microscope. The remaining plaque-derived culture was filtered through a 0.45 µm filter to remove remaining prey cells and used in a second plaque assay. To obtain pure cultures, a single derived plaque was used to start a third plaque assay. Filtrates of the final plaques were then used to generate glycerol stocks (20% (v/v)) and for subsequent DNA isolation and whole genome sequencing.

### *c) Prey cultivation*

*E. coli* strain ML35 (final OD600=1) was used as prey culture for *Bdellovibrio tomkyle* strain MYbb1, *Bdellovibrio tiberii* strain MYbb2, *Bdellovibrio kumpostii* strain MYbb5, *Bdellovibrio bagaluti* strain MYbb7 and MYbb10, *Bdellovibrio bacteriovorus* strain MYbb11, and *Bdellovampiro gaculeatus* strain SBM16. For *Bdellovibrio krueschi* strain MYbb4, *Ochrobactrum sp.* strain MYb248 (final OD600=1) was used as prey. For *Bdellovenatio daniorerio* strain ZFWA1, *Aeromonas sp.* strain AerZOR0001 ΔspdE (final OD600=1) was used as the prey culture. Prey cells ML35, MYb248, and AerZOR0001ΔspdE overnight cultures were grown in LB (Lysogeny broth) Medium or TSB (Tryptic Soy Broth), respectively, continuously shaking for 16 hours at 28°C and 180 rpm.

*d) Pseudobdellovibrionaceae cultivation*

Co-cultures of MYbb and ZFWA1 predator cells with respective prey were grown in DNB medium, incubated at 30°C, 180 rpm for 24 hours, then transferred to fresh medium with prey for another 24 hours, as described in Remy et al [4]. Before proceeding, co-cultures were filtered twice through a 0.45 µm filter to remove the remaining prey cells. For *B. gaculeatus* strain SBM16, co-culture of predator cells and *E. coli* ML35 prey cells (OD600=1) were grown in DNB medium supplemented with 2g/l sea salt, incubated at 20°C, 180 rpm for 48 hours, then transferred to fresh medium with prey for another 48 hours.

*DNA extraction and whole genome sequencing*

*a) Fish aquaculture derived isolates*

The *Pseudobdellovibrionaceae* isolates derived from fish aquaculture facilities were each grown in co-culture with the prey strain used for enrichment. Growth of the BALO strains was determined by visually monitored daily for the clearing of prey cells, and the presence of small, highly motile cells was verified with microscopy. Avoiding debris, co-cultures were transferred to a new tube and centrifuged at 10,000 x g for 2min to pellet cells. After removing the supernatant, DNA was extracted from the pelleted cells with the DNeasy® UltraClean® Microbial kit (Qiagen). Short-read sequencing was performed by SeqCenter (seqcenter.com). Briefly, libraries were prepared with the Illumina DNA Prep kit and sequenced in a 2x150bp run on an Illumina NextSeq 2000 instrument (Illumina). Long-read sequencing of *B. daniorerio* ZFWA1 was performed by Plasmidsaurus (plasmidsaurus.com) using Oxford Nanopore Technologies.

*b) Worm substrate derived isolates*

Worm substrate derived *Pseudobdellovibrionaceae* were cultivated in co-cultures with their respective prey strains as described earlier (“MYbb Cultivation”). Once a sufficient predator cell density was reached (checked via microscopy), cultures were filtered through a 0.45 μm filter and centrifuged for 10 minutes at 16,000 x g. The supernatant was removed, and the pellet was washed with 25 mM HEPES buffer. This suspension was centrifuged for 10 minutes at 16,000 x g, and the pellet was used to isolate DNA using the NucleoSpin Tissue Kit (Macherey-Nagel, Düren, Germany), following the support protocol for bacteria with minor adjustments as described in Zimmermann et al. [5]. Briefly, a RNase A digestion (0.02 mg, incubation for 20 minutes) was followed by the standard protocol for human or animal tissue and cultured cells. DNA was eluted in 50 μl pre-warmed BE buffer and used directly for sequencing without freezing. All steps were performed using Thermo Scientific™ ART™ Wide Bore Filtered Pipette Tips and DNA LoBind tubes (Eppendorf).

Library preparation for PacBio long-read sequencing was done with the SMRTbell Express Template Prep Kit 2.0 according to the manufacturer's protocol for multiplexed microbial libraries without shearing due to fragment lengths of the input DNA. Sequencing was done on a Sequel IIe, one SMRTCell 8M, Sequel Binding Kit 2.2, and Sequencing Plate 2.0. Movie time was 15 hrs. Library preparation for Illumina sequencing was done with the Illumina DNA Prep kit according to the manufacturer's protocol. Subsequent sequencing was done on the Illumina MiSeq with the MiSeq Reagent Kit v2 (500 cycles). MYbb2 was sequenced on the NextSeq 500 using the NextSeq 500/550 High Output Kit v2.5 (300 Cycles). For this, the sample was prepared with transposase Tn5 according to Picelli et al. [6].

*Microscopy*

Copper, 400 square mesh, formvar-coated TEM grids were glow discharged for 60 s at 0.6 mbar air pressure and 10 mA glow current using a Safematic CCU-010 unit, and then incubated with

10 µl of the BALO culture (dilution sample dependent) for 5-15 min. Grids were washed shortly on 7 drops of water, stained with 1% uranyl acetate for 10 s, blotted to remove the excess stain, and air dried. Samples were imaged in a Tecnai G2 Spirit BioTwin transmission electron microscope (FEI / Thermo Fisher Scientific) operated at 80 kV using TEM User interface v4.2 and equipped with a LaB6 filament. Images were recorded with an Eagle 4k x 4k CCD camera (FEI / Thermo Fisher Scientific) operated with Tia software v2.5 (all FEI / Thermo Fisher Scientific) or with a MegaView III G2 CCD camera (EMSIS), using iTEM v.5 software (both Olympus Soft Imaging solutions / EMSIS).

Size measurements of bacteria were performed on images taken at 11,000x (TIA camera) or 16,500x (MegaView III camera) magnification using FIJI software [7]. A segmented line tool was used to measure the length of flagella, and a polygon tool was used to measure the cell body area, perimeter, and maximum and minimum Feret diameter. The maximum (Max Feret) and minimum (Min Feret) Feret diameter, or caliper diameter, are defined as the longest and shortest distance between two parallel tangents on opposite sides of the cell boundary. The ratio (Max Feret / Min Feret) indicates the degree to which a particle is stretched or how similar its projected contour is to a circle. The Feret diameter measurements showed to be good indicators of the cell size and cell shape, with higher values suggesting elongated shape.

*Genome assembly and annotation*

*a) Fish aquaculture-derived isolates*

Paired-end reads obtained from Illumina NextSeq were filtered using BBDuk v37.25 (sourceforge.net/projects/bbmap/) to remove Illumina adapters (k=23, mink=11) using normal mode and to quality-trim reads to Q10. Forward and reverse paired end reads were de novo assembled with SPAdes v3.15.3 [8] with the single-cell option. Reads were aligned to the contig assembly using bowtie2 v2.4.4 [9] and converted to BAM, sorted, and indexed with samtools v1.5 [10]. Contig assemblies were then binned using MetaBAT2 v2.12.1 [11] and MaxBin2 v2.2.7 [12]. Predator contig bins were differentiated from prey contig bins and manually refined in Anvio v7.1 [13, 14]. To make the ZFWA1 hybrid assembly, Illumina reads that aligned to the refined, short-read assembly using bowtie2 were combined with long reads from Nanopore sequencing for genome assembly with SPAdes. The hybrid assembly was then manually refined in Anvio.

*b) Worm substrate derived isolates*

We obtained sequencing data for MYbb2 (NextSeq 500) and MYbb1, MYbb4, MYbb5, MYbb7, MYbb10, and MYbb11 (MiSeq + Pacbio Sequel-54349U). Raw reads from genome sequencing were filtered using fastq_illumina_filter 0.1, trimmed with PRINSEQ-lite 0.20.4 [15], and re-synchronized by repair.sh (from BBMap v38.87) [16]. The genome assembly was done with several algorithms and pipelines for comparison: i) SPAdes v3.14.1 [8], ii) MaSuRCA 3.4.2 [17], iii) Unicycler v0.4.8 (normal and bold mode) [18], iv) shovill v1.1.0 (https://github.com/tseemann/shovill), v) SKESA v2.4.0 [19]. For MYbb1, MYbb4, MYbb5, MYbb7, MYbb10, and MYbb11, additional hybrid assemblies were obtained from SPAdes, MaSuRCA, and Unicycler. Genome assembly quality was evaluated using QUAST v5.1.0rc1 [20]. For each assembly, a score was calculated based on characteristics such as the number of contigs, N50, L50, largest contig, completeness, mapped reads, properly paired reads, and N's per 100 kbp. The score was increased by 1 if the value of a characteristic belonged to the best 25% of all assemblies for an organism. The characteristics 'completeness' and 'longest contig' were considered most important and the score was increased by 2 if their value belonged to the best 5%. All assembly contigs were scrutinized for potential contamination using blobtools v1.1.1 [21]. For this means, short raw reads were sorted with samtools v1.10 [10] and mapped against the assembled contigs by BWA [22]. The assembled contigs were aligned against the nucleotide database (NCBI nt, 09/2019) employing blastn [23]. Finally, basic polishing was done based on coverage length GC plots as proposed by [24]. Contigs were filtered by length (>500bp), coverage (>5), and GC (<0.95), and the remaining contigs were considered as the final genomic sequence. Whenever possible, processes were parallelized with gnu parallel [25].

*Comparative genomic analysis*

*a) Orthogroup inference*

All complete *Pseudobdellovibrionaceae* genomes (a total of 12) were downloaded from the Bacterial and Viral Bioinformatics Resource Center (last accessed January 2023). These genomes along with the nine assembled in the present study (21 in total) were quality-checked for completeness, contamination, and strain heterogeneity using CheckM2 v1.0.2 [26]. Gene prediction and annotation on all genomes were performed with Prodigal v2.6.3 [27] wrapped in Prokka v1.14.6 [28] and eggNOG-mapper v2 [29, 30] with protein alignments with DIAMOND [31]. Ortholog prediction was performed on amino acid fasta files of *Pseudobdellovibrionaceae* genomes using OrthoFinder v2.5.4 [32] with default settings. Orthologs were plotted using the UpSetR v1.4.0 package [33] in R v4.2.2 (R Core Team, 2022).

*b) Phylogenomic analysis of* Pseudobdellovibrionaceae

To determine the phylogenetic relationships between the *Pseudobdellovibrionaceae* strains, DNA sequences of single-copy orthologous genes (orthogroups with a single gene member) common to all genomes were used to generate a phylogenomic tree. The nucleotide sequences of each single-copy orthogroup from each *Pseudobdellovibrionaceae* genome were aligned using MAFFT v4.475 [34]. Genes were concatenated by genome, and a tree was constructed using IQ-TREE v2.1.2 [35] with 1000 bootstrap replicates using UFBoot2 [36]. Within IQ-TREE, ModelFinder [37] determined the most likely model was the general time reversible model with empirical base and codon frequencies, allowing for a proportion of invariable sites, and a discrete Gamma model with default four rate categories (GTR+F+I+G4).

### *c) Phylogenomic analysis of prokaryotes*

Similarly, a species tree was constructed to determine the phylogenetic relationships between various predatory and non-predatory prokaryotes. Representative genomes were included to span distant and close relatedness to Bdellovibrionata as well as a variety of predatory and non-predatory bacteria based on the revised 16S phylogeny from Kamada et al. [38]. Ortholog prediction was performed on amino acid fasta files of prokaryotic genomes using OrthoFinder v2.5.4 [32] with the *-msa* option for multiple sequence alignment using MAFFT [34] as the default aligner. The amino acid species tree multiple sequence alignment was then used as input into IQ-TREE [35] with 1000 bootstrap replicates using UFBoot2 [36]. Within IQ-TREE, ModelFinder [37] determined the most likely model was the general matrix model with empirical base and codon frequencies, allowing for a proportion of invariable sites, and a discrete Gamma model with default four rate categories (LG+F+I+G4). Trees were rooted using the Interactive Tree of Life v6 [39] and visualized with the ggtree v3.12.0 [40] and treeio v1.28.0 [41] packages in R.

### *d)* Pseudobdellovibrionaceae *species identification*

Taxonomic affiliation was assigned using 95% average nucleotide identity (ANI) calculated with fastANI v1.34 and 65% average amino acid identity (AAI) calculated with CompareM v0.1.2 (https://github.com/dparks1134/CompareM) as suggested by Jain et al. and Konstantinidis et al., respectively [42,43].

### *Analysis of* Pseudobdellovibrionaceae *prevalence in host microbiome data*

16S rDNA sequences of all *Pseudobdellovibrionaceae* strains used in this study were combined to create a database using the makeblastdb function in blast [23]. 16S rDNA amplicon datasets from studies of zebrafish, stickleback, and *C. elegans* microbiomes were then screened for the presence of our *Pseudobdellovibrionaceae* strains using blastn, allowing for up to 4 mismatches (i.e. at least 99% identity). *Pseudodellovibrionaceae* ASVs appearing exclusively in negative controls were excluded. ASVs that appeared in both host samples and negative controls were permitted when their relative abundance in host samples exceeded their relative abundance in negative controls by 2.5 times.

**Table S1. Size measurements for *Pseudobdellovibrionaceae* strains described in this study.** Measurements of bacteria were performed on images taken at 11,000x (TIA camera) or 16,500x (MegaView III camera) magnification using FIJI software (Schindelin et al., 2012). A segmented line tool was used to measure the length of flagella, and a polygon tool was used to measure the cell body area, perimeter and Feret diameter. Culture column indicates whether fresh prey was added before imaging or whether standard overnight cultures were used. The mean, standard deviation (SD) and count numbers are displayed for flagellum and cell body measurements. Units for each measurement are indicated in square brackets. Count column refers to the number of images analyzed for mean values. Run column indicates how many independent experiments (the first number) and how many independent grids (the second number) were analyzed for each sample.

| | | | | **Flagellum** [nm] | | | **Cell body** [nm2] | | | [nm] | | [nm] | | [nm] | | **Max/Min Feret** |
|---|---|---|---|---|---|---|---|---|---|---|---|---|---|---|---|---|
| | | | | **Length** | | | **Area** | | | **Perimeter** | | **Max Feret** | | **Min Feret** | | |
| **Run** | **Culture** | **Strain** | **Species** | Mean | SD | Count | Mean | SD | Count | Mean | SD | Mean | SD | Mean | SD | |
| **Exp-1-1** | **fresh** | **MYbb11** | ***Bdellovibrio bacteriovorus*** | 2869.91 | 571.00 | 23 | 218367.18 | 29477.59 | 28 | 1853.18 | 165.43 | 717.00 | 72.76 | 395.55 | 28.90 | 1.81 |
| **Exp-1-1** | **standard** | **MYbb2** | ***Bdellovibrio tiberii*** | 3527.38 | 364.46 | 38 | 217236.98 | 37231.96 | 37 | 1914.69 | 170.86 | 742.84 | 69.89 | 393.59 | 42.40 | 1.89 |
| **Exp-2-1** | **standard** | **MYbb2** | ***Bdellovibrio tiberii*** | 3293.90 | 271.87 | 13 | 216840.47 | 26979.45 | 14 | 1835.21 | 155.91 | 715.66 | 70.25 | 386.04 | 30.11 | 1.85 |
| **Exp-2-2** | **fresh** | **MYbb2** | ***Bdellovibrio tiberii*** | 3126.97 | 342.25 | 5 | 333513.61 | ######## | 5 | 2319.42 | 485.76 | 930.85 | 214.59 | 454.64 | 57.20 | 2.05 |
| **Exp-2-3** | **fresh** | **MYbb2** | ***Bdellovibrio tiberii*** | 3139.95 | 418.63 | 7 | 299488.35 | 44740.77 | 7 | 2200.10 | 186.41 | 864.96 | 80.04 | 447.99 | 26.71 | 1.93 |
| **Exp-3-1** | **standard** | **MYbb2** | ***Bdellovibrio tiberii*** | 3619.76 | 295.27 | 23 | 198697.24 | 28957.71 | 24 | 1818.49 | 159.84 | 710.99 | 67.91 | 371.14 | 31.98 | 1.92 |
| **Exp-3-2** | **fresh** | **MYbb2** | ***Bdellovibrio tiberii*** | 3308.36 | 456.41 | 6 | 244338.78 | 11805.88 | 6 | 1992.26 | 64.09 | 774.40 | 34.33 | 419.35 | 19.61 | 1.85 |
| **Exp-1-1** | **standard** | **MYbb5** | ***Bdellovibrio kumpostii*** | 3015.21 | 552.77 | 37 | 211612.05 | 27309.06 | 43 | 1900.93 | 164.99 | 765.42 | 78.32 | 368.15 | 30.88 | 2.08 |
| **Exp-1-1** | **fresh** | **MYbb1** | ***Bdellovibrio tomkyle*** | 2927.00 | 205.00 | 25 | 182847.10 | 20264.22 | 29 | 1642.49 | 99.33 | 636.82 | 47.12 | 364.51 | 32.34 | 1.75 |
| **Exp-1-1** | **standard** | **MYbb1** | ***Bdellovibrio tomkyle*** | 3119.83 | 242.85 | 10 | 188129.36 | 18184.82 | 10 | 1692.69 | 97.61 | 662.40 | 48.83 | 366.91 | 19.10 | 1.81 |
| **Exp-1-1** | **standard** | **MYbb7** | ***Bdellovibrio bagaluti*** | 3029.98 | 430.80 | 21 | 190591.01 | 30690.53 | 25 | 1782.26 | 178.05 | 708.43 | 83.62 | 356.03 | 26.66 | 1.99 |
| **Exp-1-2** | **standard** | **MYbb7** | ***Bdellovibrio bagaluti*** | 2945.00 | 448.43 | 39 | 219423.04 | 31309.96 | 40 | 1854.31 | 205.64 | 741.26 | 102.39 | 364.54 | 21.95 | 2.03 |
| **Exp-1-1** | **standard** | **MYbb10** | ***Bdellovibrio bagaluti*** | 2898.59 | 344.31 | 29 | 286414.85 | 36332.68 | 73 | 2364.92 | 213.95 | 966.73 | 97.28 | 422.27 | 33.72 | 2.29 |
| **Exp-1-1** | **standard** | **MYbb4** | ***Bdellovibrio krueschi*** | 2562.57 | 314.55 | 32 | 206757.73 | 30450.69 | 37 | 1974.34 | 197.71 | 795.06 | 96.20 | 365.80 | 27.10 | 2.17 |
| **Exp-2-1** | **standard** | **MYbb4** | ***Bdellovibrio krueschi*** | 2598.53 | 339.85 | 23 | 289570.53 | 38234.62 | 37 | 2326.73 | 194.58 | 937.27 | 97.14 | 436.66 | 40.56 | 2.15 |
| **Exp-1-1** | **standard** | **ZFWA1** | ***Bdellovenatio daniorerio*** | 2684.55 | 387.96 | 37 | 334917.86 | 56471.14 | 36 | 2299.92 | 224.09 | 897.21 | 96.86 | 480.81 | 36.29 | 1.87 |
| **Exp-1-1** | **standard** | **ZFWA1** | ***Bdellovenatio daniorerio*** | 2790.70 | 292.89 | 16 | 246342.07 | 81129.19 | 12 | 2086.76 | 168.85 | 818.36 | 79.63 | 431.31 | 40.87 | 1.90 |
| **Exp-2-1** | **standard** | **ZFWA1** | ***Bdellovenatio daniorerio*** | 2846.00 | 202.56 | 43 | 207896.05 | 28765.22 | 56 | 1804.42 | 150.77 | 703.60 | 66.46 | 382.45 | 28.44 | 1.84 |

| | | | | | | | | | | | | | | | | |
|---|---|---|---|---|---|---|---|---|---|---|---|---|---|---|---|---|
| **Exp-2-2** | **standard** | **ZFWA1** | ***Bdellovenatio daniorerio*** | 2774.20 | 137.27 | 8 | 212479.19 | 36445.40 | 7 | 1863.77 | 200.89 | 736.40 | 94.40 | 374.77 | 23.06 | 1.96 |
| **Exp-3-1** | **standard** | **ZFWA1** | ***Bdellovenatio daniorerio*** | 2737.65 | 226.21 | 11 | 235290.08 | 49711.29 | 16 | 1919.09 | 260.78 | 758.66 | 118.00 | 392.95 | 37.93 | 1.93 |
| **Exp-1-1** | **standard** | **SBM 16** | ***Bdellovampiro gaculeatus*** | 7128.74 | 1573.09 | 20 | 772561.00 | 236087 | 21 | 4328.00 | 968.00 | 1886.00 | 459.00 | 582.00 | 77.00 | 3.24 |

**Table S3. Genbank accession numbers of genome assemblies included in phylogenomic analyses.** Prokaryotic genomes were used to infer phylogenetic relationships between periplasmic and epibiotic BALOs. *Pseudobdellovibrionaceae* genomes were used to infer phylogenetic relationships between periplasmic and epibiotic existing and novel *Pseudobdellovibrionaceae* strains.

| **Species** | **Strain** | **Genbank Accession** |
|---|---|---|
| Vampirovibrio chlorellavorus | Vc_AZ_1 | GCA_003149375.1 |
| Micavibrio aeruginosavorus ARL-13 | ARL-13 | GCA_000226315.1 |
| Stenotrophomonas maltophilia | NCTC10257 | GCA_900186865.1 |
| Myxococcus xanthus DK 1622 | DK 1622 | GCA_000012685.1 |
| Bacteriovorax stolpii | DSM 12778 | GCA_002872415.1 |
| Halobacteriovorax marinus SJ | SJ | GCA_000210915.2 |
| Peredibacter starrii | A3.12 | GCA_034259205.1 |
| Pseudobacteriovorax antillogorgiicola | DSM 103413 | GCA_004346105.1 |
| Ensifer adhaerens | Casida A | GCA_000697965.2 |
| Oligoflexus tunisiensis | Shr3 | GCA_001748245.1 |
| Nitrospina gracilis | 3/211 | GCA_000341545.2 |
| Geobacter sulfurreducens | PCA | GCA_000007985.2 |
| Desulfovibrio vulgaris | DP4 | GCA_000015485.1 |
| Acidobacterium camsulatum | ATCC 51196 | GCA_000022565.1 |
| Nanoarchaeum equitans | Kin4-M | GCA_000008085.1 |
| *Bdellovibrio/Pseudobdellovibrio* | | |
| Bdellovibrio bacteriovorus str. Tiberius | Tiberius | CP002930.1 |
| (Pseudo)-Bdellovibrio exovorus JSS | JSS | CP003537.1 |
| Bdellovibrio sp. qaytius | qaytius | CP025734 |
| Bdellovibrio sp. NC01 | NC01 | CP030034 |
| Bdellovibrio sp. ZAP7 | ZAP7 | CP030082 |
| Bdellovibrio bacteriovorus HD100 | HD100 | BX842601 |
| Bdellovibrio sp. KM01 | KM01 | CP058348 |
| Bdellovibrio sp. LBG001 | LBG001 | CP093442 |
| Bdellovibrio bacteriovorus W | W | CP002190.1 |
| Bdellovibrio bacteriovorus strain SSB218315 | SSB218315 | CP020946 |
| Bdellovibrio bacteriovorus strain 109J | 109J | CP007656.1 |
| Bdellovibrio bacteriovorus kdesi | kdesi | CP102930 |

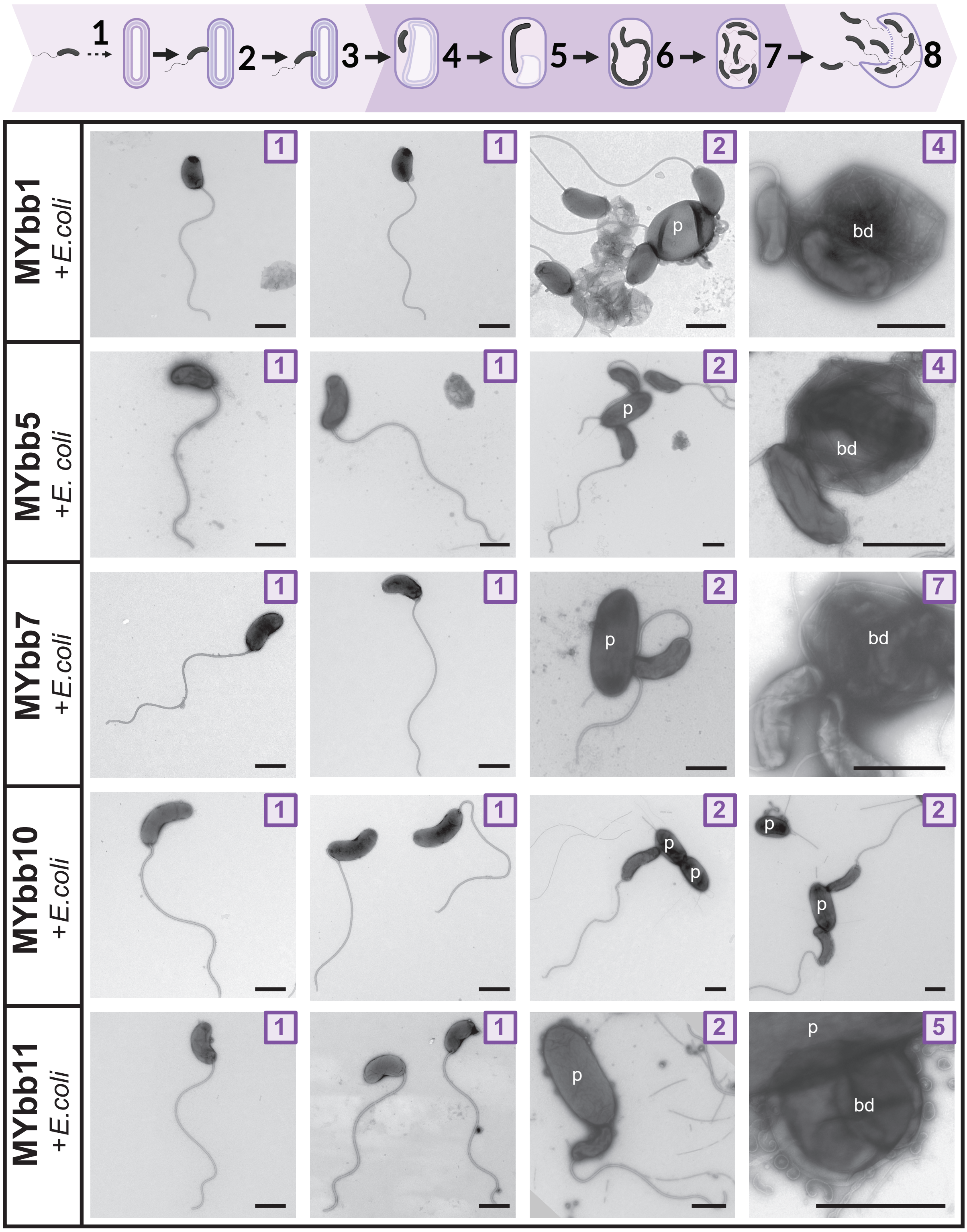

**Figure S1: Cell shape and predation mode of periplasmic *Pseudobdellovibrionaceae* strains MYbb1, MYbb5, MYbb7, MYbb10 and MYbb11.** Stages in the life cycle of periplasmic *Pseudobdellovibrionaceae* isolates are shown on top. Numbers in image corners correspond to the proposed life cycle stage. Stage 1 images represent *Pseudobdellovibrionaceae* cells in attack phase. In stage 2, predators are attached to prey cells (p). After penetration of the prey's outer membrane, *Pseudobdellovibrionaceae* cells enter stages 4-7 with intraperiplasmic growth in bdelloplasts (bd), with one or more predator cells confined within a prey cell (p). Samples were imaged by TEM after negative staining.

Scale bar is 500 nm. Life cycle images created in BioRender. Wülbern, J. (2024) BioRender.com/t31h353.

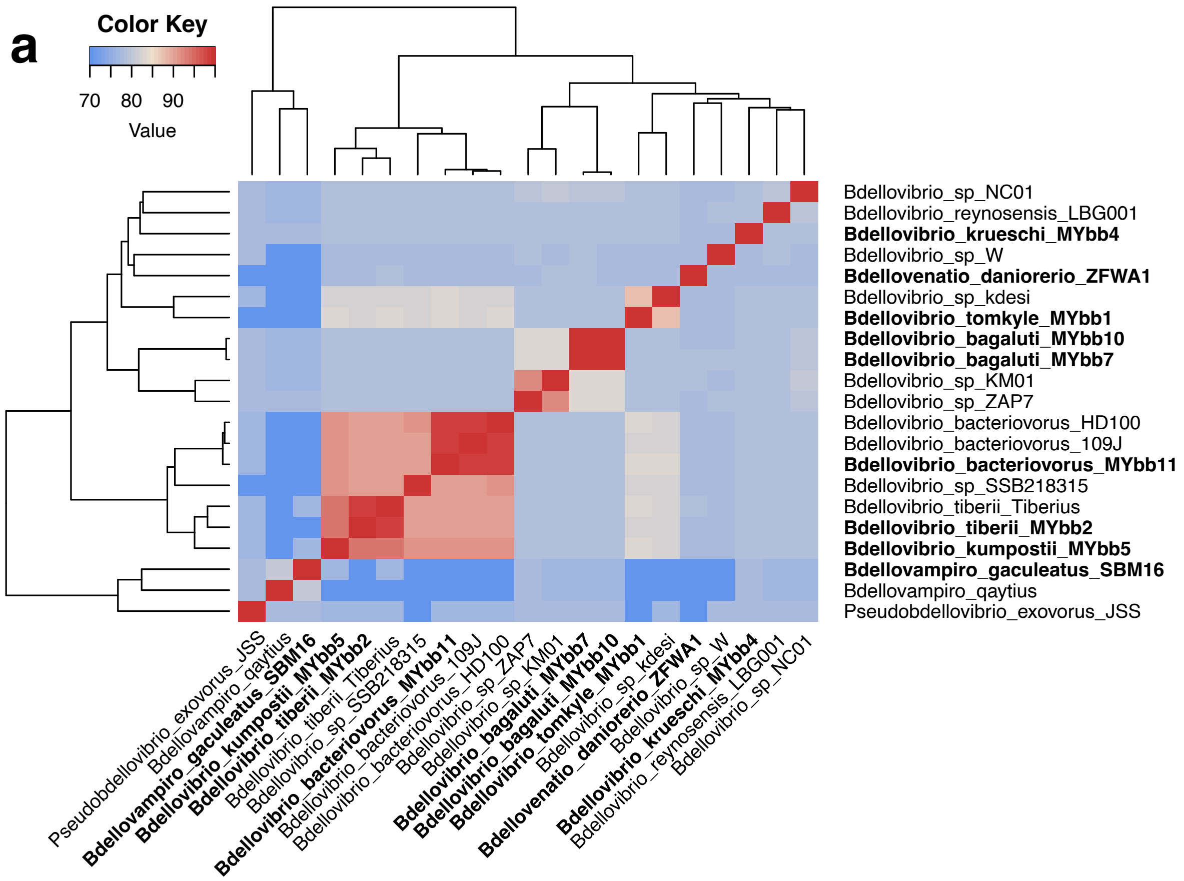

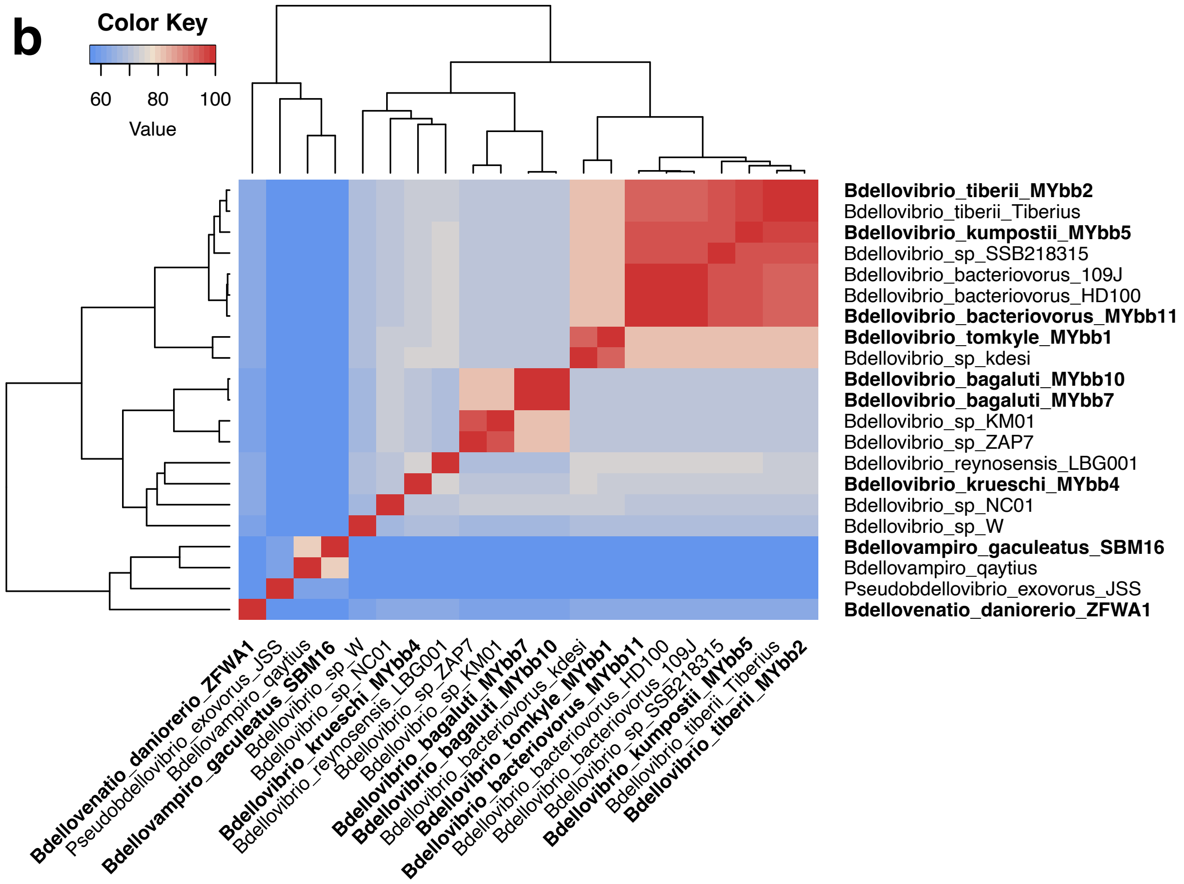

**Figure S2. Average Nucleotide Identity (a) and Average Amino Acid Identity (b) between strains.** Pairwise ANI was calculated with fastANI v1.34 and pairwise AAI was calculated with CompareM v0.1.2. Both were visualized in R with ComplexHeatmap v2.20.0. Novel isolates are listed in bold.

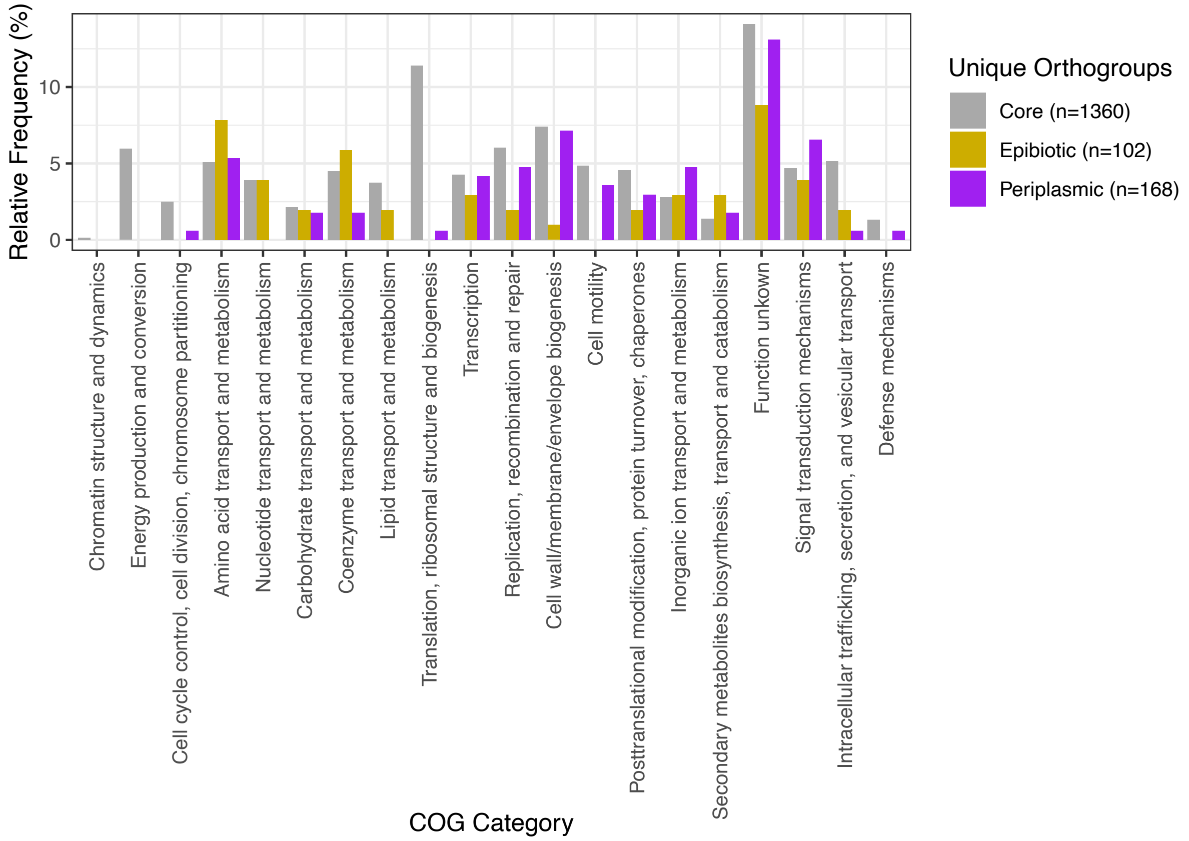

**Figure S3. Relative frequency of COG categories in orthogroups unique to the core, epibiotic, and periplasmic genomes.** Orthogroups were annotated with a COG category using eggNOG-mapper and classified as core (found in all genomes), epibiotic, or periplasmic. Total orthogroups per classification are listed in the figure legend. Relative frequency is calculated by summing COG annotations per category within a classification and divided by the total number of orthogroups per classification.
